# A Cross-Species Systems Genetics Framework Identifies Causal Genes in Diabetic Nephropathy

**DOI:** 10.64898/2026.06.30.735464

**Authors:** Kunal Mishra, Rashidah Binte Sakban, Sujithra Shankar, Jasmine Wong, Benjamin L. Farah, Jing Guo, Jianhong Ching, Ming Shen Tham, Jean-Paul Kovalik, Susan B. Gurley, Enrico Petretto, Nicholas Stanislaw Tolwinski, Thomas M. Coffman, Jacques Behmoaras

## Abstract

Diabetic nephropathy (DN) is the leading cause of kidney failure in the developed world, but the genetic architecture of DN susceptibility is not well characterised. Here we apply a systems genetics approach in a mouse model of DN to discover novel QTLs for clinically relevant phenotypes including albuminuria, glomerulosclerosis, and macrophage infiltration. For context and prioritisation, we combined single-cell-transcriptomics-guided pQTL and eQTL mapping with cell-type-specific co-expression networks, identifying 192 candidate pGenes for albuminuria. While many were novel, 27% had prior genetic associations, and 40% were validated in a human diabetic cohort. Twelve genes belong to a podocyte network enriched for human GWAS signals. Among those, functional significance of the E3 ubiquitin ligases *DCAF6* and *ZNRF2* was confirmed by knockdown in Drosophila nephrocytes. Our systems genetics approach identified DN susceptibility genes previously validated in human GWAS while uncovering potential new genes and pathways that could be exploited for risk stratification and therapeutics development.

## Introduction

Diabetic nephropathy (DN) is among the most common causes of end-stage kidney disease (ESKD) worldwide, arising as a complication of diabetes mellitus^1–4^. Only 30-40% of individuals with diabetes develop DN^5,6^, and familial clustering indicates that genetic background is a major determinant of DN susceptibility^7–9^. The precise molecular determinants of this susceptibility remain incompletely defined. Although multiple large-scale GWAS have been conducted in human cohorts with diabetes enriched for complications, the genetic architecture of DN susceptibility remains poorly resolved, with few robustly replicated loci and limited mechanistic insight into causal genes^10,11^. This likely reflects the phenotypic heterogeneity of diabetic kidney disease across human populations, as well as the fundamental challenge of translating statistical associations into functionally validated pathogenic mechanisms.

Albuminuria is a sensitive and clinically powerful biomarker of glomerular injury that predicts progressive kidney disease and cardiovascular risk in individuals with diabetes^12,13^. It emerges early in the course of DN, reflects disruption of the glomerular filtration barrier more directly than composite markers of kidney function, and stratifies individuals at greatest risk of progression to ESKD. Using albuminuria as the primary mapping trait in genetic studies, therefore, provides a disease-proximal and mechanistically informative quantitative phenotype for dissecting the genetic architecture of DN susceptibility.

Despite large-scale GWAS of urinary albumin-to-creatinine ratio in both general and diabetic populations, the number of robustly identified loci remains modest relative to the heritability of albuminuria, and the causal mechanisms underlying most signals remain undefined^14,15^. Animal models can offer a powerful complementary strategy to overcome limitations of human genetic studies^16–18^. Crosses using inbred mouse lines can provide simplified and controlled genetic backgrounds, longitudinal phenotyping, and direct assessment of target organ pathology. Quantitative trait loci (QTL) mapping in genetically segregating populations allows systematic linkage of genetic variation to continuous disease traits such as albuminuria, kidney pathology, and immune cell infiltration^19–23^. Integrating molecular, metabolic and cellular profiling, a systems genetics approach^24^ provides a framework for prioritising and testing candidate genes, which is often challenging in human cohorts. Critically, candidate genes emerging from systems genetics studies can be directly tested for association in large human cohorts, providing a direct route from mechanistic discovery to translational relevance.

Historically, the development of animal models of diabetic complications, including nephropathy, has been challenging, and it has been difficult to create models that fully recapitulate the human disorder^18,25^. Nonetheless, we have previously created a mouse model with many characteristic features of human DN, including hyperglycaemia, marked albuminuria, glomerulosclerosis, renal inflammation, and hypertension. Moreover, this model exhibits dramatic strain-dependent differences in DN susceptibility. On the susceptible 129/SvEv background, mice develop florid signs of DN, whereas the C57BL/6 strain is resistant^26–28^. This strain-dependent susceptibility, in a defined diabetic and hypertensive background, provides a tractable genetic system for mapping molecular determinants of early DN independently of glycaemic confounding. Here, we leverage this model to develop and validate a multi-layered systems genetics framework that integrates mouse single-cell transcriptomics-guided QTL mapping, eQTL mapping, cell type-specific gene co-expression network analysis, invertebrate functional validation, and genetic association in diabetic humans to systematically prioritise and characterise candidate genes underlying albuminuria and kidney injury in DN.

## Results

### Progressive albuminuria and kidney damage in F2 mice

To systematically dissect the genetic and cellular determinants of diabetic nephropathy, we designed a multi-modal F_2_ QTL mapping study integrating genetics, single cell and bulk transcriptomics, phenotyping, metabolomics, human cohort validation, and functional validation of candidate genes in *Drosophila* (Figure 1A). 129-Akita/ReninTg (129AR) and C57BL/6NJ-Akita/ReninTg (BL6AR) mice were bred and maintained as described in the methods. As previously reported^28^, 24-week-old 129AR mice develop albuminuria, glomerulosclerosis, tubular thickening, and marked macrophage infiltration, evidenced by CD68+ deposits in the renal cortex (Figure 1B). In contrast, age-matched BL6AR mice did not exhibit these pathological features, despite comparable serum glucose levels and the presence of both the Akita mutation and the Renin transgene (Figure 1B, C). We exploited this strain-dependent susceptibility to diabetic nephropathy to generate an F2 cross population (n=279), enabling systematic mapping of phenotypic variability and identification of genetic determinants of disease. F2 mice were aged to 24 weeks, with glucose levels and albuminuria measured every 8 weeks; DNA was collected for whole-genome sequencing (WGS), kidneys were harvested for total RNA isolation, histology, and IHC, and urine was collected for targeted organic acid metabolomics (Figure 1A).

**Figure 1.**
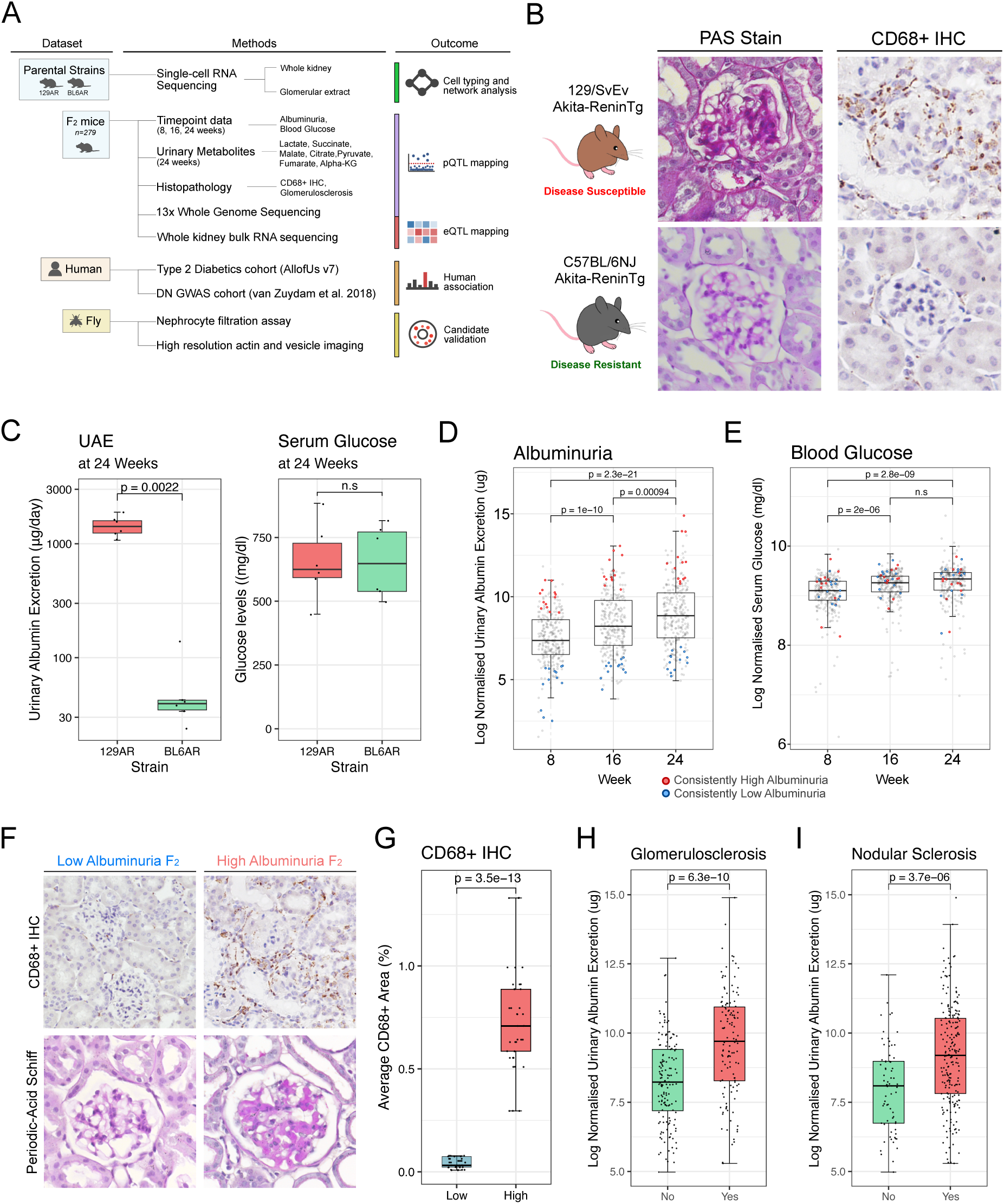
F2 mice display broad phenotypic variation across diabetic nephropathy traits. Schematic overview of the F2 QTL mapping study design, including data modalities collected from parental strains and F2 mice, human validation cohorts, and Drosophila functional validation. Boxes are color-coded by analysis type: eQTL, pQTL, cell typing, association, and validation. (B) Representative Periodic Acid-Schiff (PAS) histology and CD68 immunohistochemistry (IHC) images from 129/SvEv Akita-ReninTg (129AR, disease susceptible) and C57BL/6NJ Akita-ReninTg (BL6AR, disease resistant) mice. (C) Urinary albumin excretion (UAE) and serum glucose levels in 129AR and BL6AR mice (n=6 per group). Significance was assessed using the Wilcoxon rank-sum test. (D, E) Boxplots showing albuminuria (D) and blood glucose (E) across the F2 population at each time point. Mice consistently ranked within the high-and low-albuminuria groups, as defined in the Methods, are highlighted in red and blue, respectively. Significance was assessed using Student’s t-test. (F) Representative PAS histology and CD68 IHC images from F2 mice in the low- and high-albuminuria groups. (G) Quantification of CD68+ staining area in the low- and high-albuminuria groups. Significance was assessed using the Wilcoxon rank-sum test. (H, I) Boxplots showing normalised UAE levels in F2 mice stratified by presence or absence of glomerulosclerosis (H) and nodular sclerosis (I). Significance was assessed using Student’s t-test.

F2 mice exhibited a broad range of albuminuria (Figure 1D, Suppl. Table 2), reflecting significant inter-individual variability, and showed a progressive increase in albuminuria over time. Blood glucose levels rose between weeks 8 and 16, with no significant increase thereafter (Figure 1E). Mice at the extremes of albuminuria levels (designated as High or Low albuminuria groups) displayed clear differences in renal histology, including the degree of sclerosis and macrophage infiltration (Figure 1F, G). Differential RNA expression between the two groups revealed distinct pathway signatures. Mice with high albuminuria showed upregulation of canonical pathogenic pathways, including inflammatory responses, epithelial-mesenchymal transition, and TNF-alpha signalling (Extended Figure 1A, B, C). Conversely, mice in the low albuminuria group upregulated genes associated with metabolic pathways, vascular transport, and fatty acid metabolism, which are processes linked to renal protection in DN^29,30^. Consistent with prior reports^31^, several urinary metabolites correlated with albuminuria levels, including lactate, succinate, pyruvate, and citrate levels (Suppl. Figure 1). Mice with more severe nodular and glomerular sclerosis also exhibited higher albuminuria (Figure 1H, I), despite having comparable blood glucose levels (Suppl. Figure 2). These findings establish the F2 population as a phenotypically heterogeneous resource with clear genetic and molecular stratification, enabling systematic mapping of albuminuria and kidney injury (macrophage infiltration and glomerulosclerosis) independently of blood glucose levels.

### QTL mapping and integrative scRNA-seq in DKD

We conducted QTL mapping for all measured traits, including albuminuria, glomerulosclerosis, macrophage infiltration, and urinary metabolites, using linear mixed models. Multivariate QTL mapping of albuminuria identified 28 significant phenotypic QTLs (pQTLs). Two loci overlapped with chromosomal regions previously associated with albuminuria in crosses between damage-susceptible SM/J and non-susceptible MRL/MpJ mice^32^, confirming the veracity of our approach. Together, these 28 pQTLs encompassed 2,325 protein-coding genes. Since not all genes within a locus are expected to be causative, we applied additional filtering to prioritise putative disease genes. Genes were retained if (i) they had human orthologs, (ii) harboured sequence differences between the parental strains and (iii) were expressed in the mouse kidney. This approach reduced the list to 192 albuminuria-associated phenotypic QTL genes (pGenes, Suppl. Table 4). To provide cellular context for downstream genetic analyses, we also performed single-cell sequencing on parental AR and WT strains, using freshly isolated glomerular and whole-kidney cells from 10-week-old mice (Figure 1A, 2A–C, Suppl. Figures 3, 4). At the single-cell level, albuminuria pGenes were predominantly expressed in podocytes and largely downregulated in the DN-susceptible 129 strain, suggesting that reduced podocyte gene activity is a major contributor to disease susceptibility (Figure 2E). Notably, of the 192 albuminuria-associated pGenes, 52 genes had been previously linked to DN-related traits in humans, including CKD, T1D, T2D, eGFR decline, and hypertension (Suppl. Table 15, Extended Figure 2), representing a significant enrichment (hypergeometric p = 7.93 x 10^-33^). Furthermore, we identified 4 genes (GNG2, KAZN, OXR1, PTK2) that have been previously associated with DN in a Finnish population^33^. The same framework identified 5 pQTLs (25 pGenes) for glomerulosclerosis (Figure 2H). Glomerulosclerosis pGenes showed patterns similar to the albuminuria signals, including predominant podocyte expression with downregulation in the 129AR strain (Figure 2I). In addition, we identified a glomerulosclerosis-associated pQTL on chromosome 4 overlapping the AGTRAP-PLOD1 locus, which has previously been implicated in hypertension and renal damage^34,35^.

**Figure 2.**
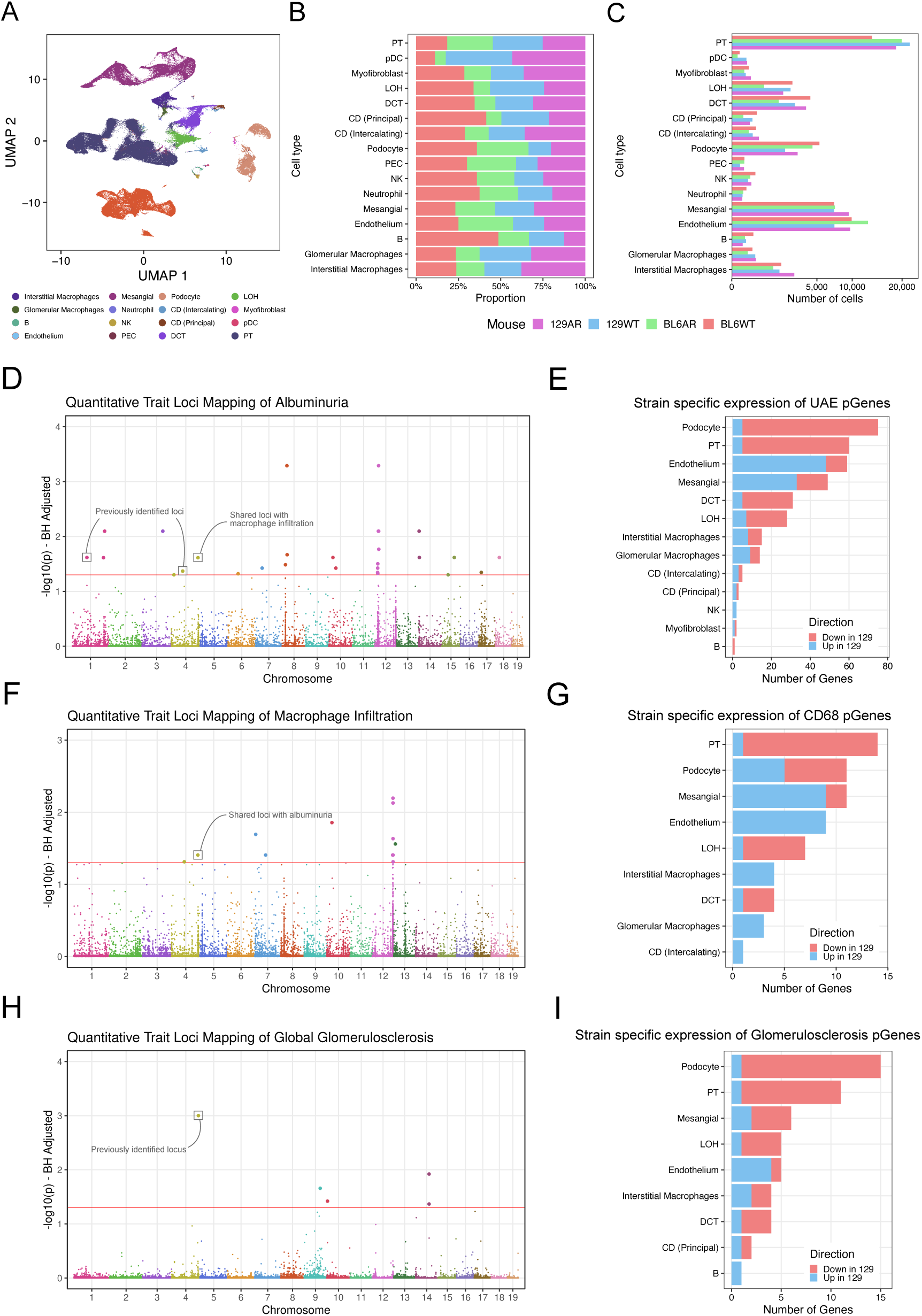
Single-cell profiling and genomic association of diabetic nephropathy traits. (A) Uniform Manifold Approximation and Projection (UMAP) showing kidney cell types captured across all strains. (B, C) Bar plots showing the proportion (B) and total number (C) of cells per strain. (D, F, H) Manhattan plots showing QTL mapping results for albuminuria (D), macrophage infiltration (F), and glomerulosclerosis (H). Horizontal lines indicate the significance threshold after multiple testing correction. (E, G, I) Bar plots showing the number of pGenes with strain-specific differential expression across kidney cell types, for albuminuria pGenes (E), macrophage infiltration pGenes (G), and glomerulosclerosis pGenes (I). Cell types are shown on the y-axis and bars are coloured by direction of expression relative to the 129 strain (red, lower in 129; blue, higher in 129).

We identified 19 pQTLs (36 pGenes) and 5 pQTLs (25 pGenes) for macrophage infiltration and glomerulosclerosis, respectively (Figure 2F-H; Suppl. Table 5, 6, 7, 8). In contrast to albuminuria pGenes, which were predominantly expressed in podocytes, macrophage infiltration pGenes were most highly expressed in proximal tubule cells and largely downregulated in the 129AR strain (Figure 2G), suggesting tubular injury as a driver of immune cell recruitment in this model. Furthermore, 4 pQTLs (12 pGenes) and 13 pQTLs (27 pGenes) were identified for urinary succinate and malate levels, respectively (Suppl. Table 9, 10, 11, 12; Extended Figure 3A-D).

In parallel, we performed eQTL mapping in the F2 cohort to further refine candidate genes within pQTL intervals (Extended Table 1; Suppl. Tables 13 and 14). We identified 5,494 *cis*-regulated genes and 50 *trans*-regulated genes, including one *trans* hotspot at 15:19836442 within a 14.8 Mb interval that encompasses 137 genes, potentially regulating 42 genes. Among the 192 albuminuria-linked pGenes, 82 were *cis*-regulated (42%), and 26 were down-regulated in podocytes, further supporting their prioritisation as putative causal drivers of albuminuria in DN. Interestingly, one pQTL on chromosome 4 (4:144879108) was shared between macrophage infiltration and albuminuria, suggesting a common genetic regulatory mechanism underlying both traits.

In summary, this integrated scRNA-seq, pQTL and eQTL mapping identified novel albuminuria-associated podocyte genes in DN. We next sought to focus on the 192 albuminuria pGenes and investigate their relevance to human DN in mechanistic contexts facilitated by systems genetics approaches.

### Disease-relevant cell-specific networks in diabetic nephropathy

To assess relevance to human kidney disease, we first examined DNA sequence variants across all albuminuria genes for association with the urine albumin-to-creatinine ratio (UACR) in diabetic participants of the All of Us Research Program (The All of Us Research Program Investigators, 2019), an independent, large-scale human cohort. After adjusting for age, sex, ethnicity, and fasting glucose, 72 of the 179 human orthologous albuminuria pGenes (40.2%) reached empirical FDR significance and 55 (30.7%) showed nominal significance (Figure 3), confirming that mouse-derived albuminuria candidates are enriched for UACR-associated variants in humans. Genes reaching FDR or nominal significance also harboured more high-confidence (HC) loss-of-function (LoF) variants per gene and a higher proportion of LoF variants than non-significant pGenes (Extended Figure 4), further supporting the functional relevance of these loci to renal albumin excretion. Overall, these results establish the human translational relevance of this pGene set.

**Figure 3.**
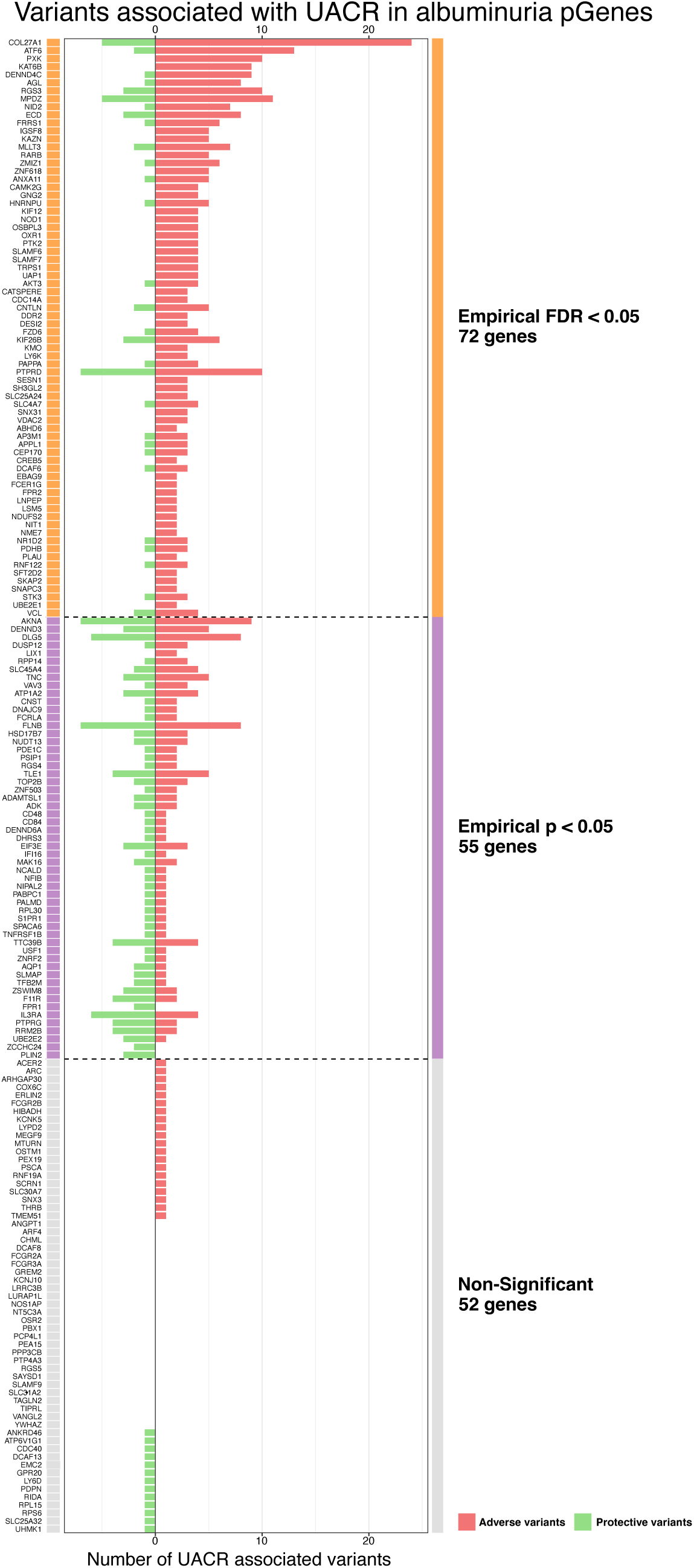
Albuminuria pGenes are enriched for UACR-associated variants in diabetic individuals. Bar plot showing the number of adverse (red) and protective (green) UACR-associated variants per albuminuria pGene, tested in 3,553 diabetic participants of the All of Us Research Program. Genes are ordered and grouped by significance tier: empirical FDR < 0.05 (72 genes), empirical p < 0.05 (55 genes), and non-significant (52 genes), separated by dashed vertical lines. A colour-coded strip beneath gene names indicates chromosomal origin.

To further annotate candidates with both human genetic support and mechanistic context, we integrated the UACR association results with cell-type-specific coexpression network analysis, combining two complementary annotation strategies to converge on a tractable set of functional candidates (Figure 4A). We thus constructed 95 cell-type-specific gene coexpression networks from parental strain single-cell data and annotated them for disease relevance (Suppl. Table 16). For each network, we examined if (i) the network eigengene correlated with albuminuria at any time point, (ii) the network genes contained any albuminuria pGenes identified through the pQTL mapping and prioritisation, and (iii) if the network genes had SNPs enriched in a human DN GWAS stratified by albuminuria^36^. This approach characterised several modules across different renal cell types (Suppl. Table 16). Given the predominant podocyte expression of albuminuria pGenes (Figure 2E), we focused on characterising networks expressed in podocytes (Figure 4).

**Figure 4.**
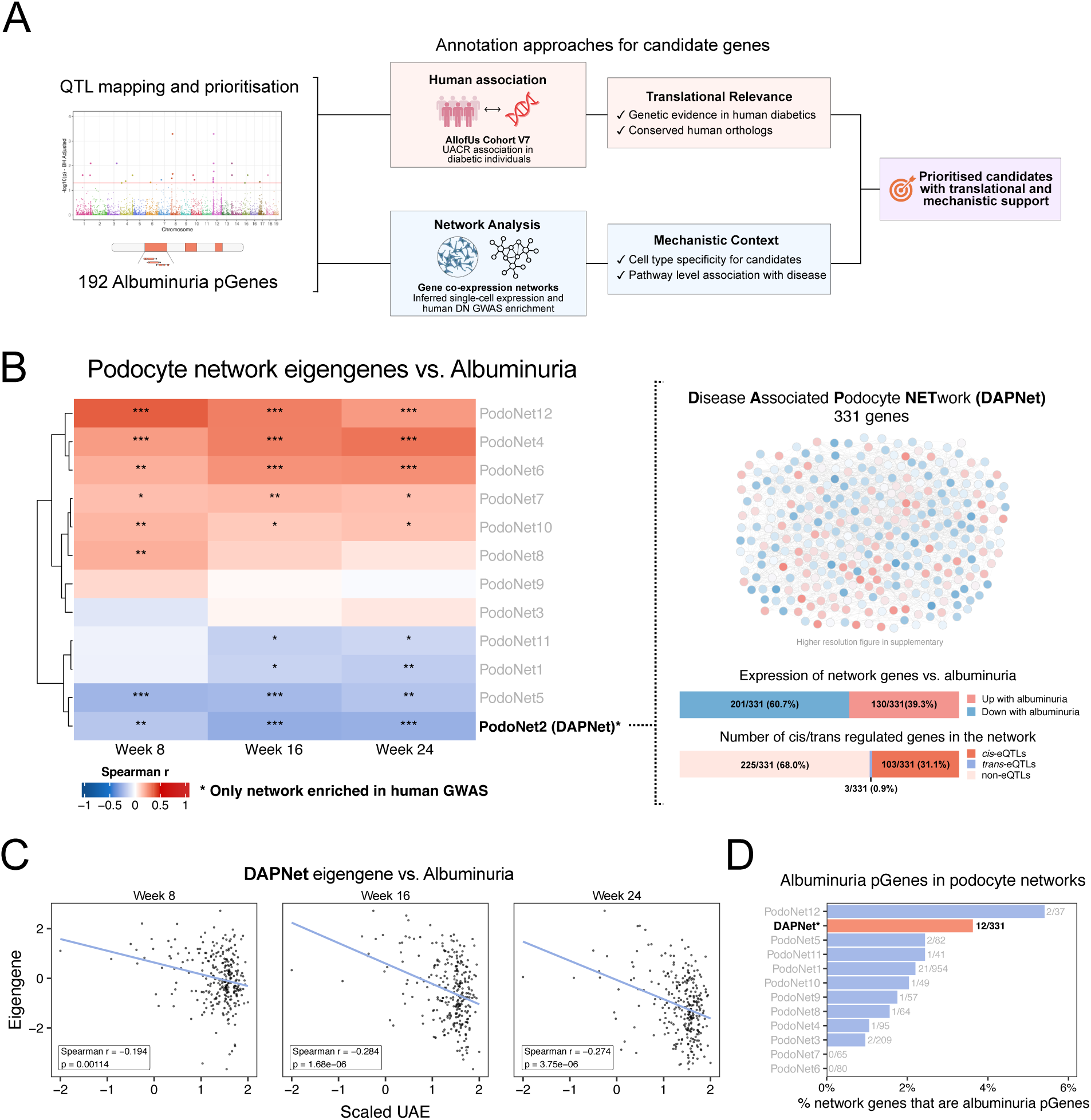
Cell-type-specific network analysis identifies the Disease-Associated Podocyte Network (DAPNet) as a disease-relevant gene module in diabetic nephropathy. (A) Schematic overview of the two complementary annotation strategies used to prioritise albuminuria candidate genes: human variant association in the All of Us diabetic cohort and cell-type-specific gene coexpression network analysis. Both approaches converge on candidates with translational relevance and mechanistic context. (B) Left: Heatmap showing Spearman correlation between podocyte network eigengenes and albuminuria at weeks 8, 16, and 24 in the F2 cohort. Asterisks indicate significance after Benjamini-Hochberg correction (* p < 0.05, ** p < 0.01, *** p < 0.001). PodoNet2 (DAPNet), the only network enriched in human DN GWAS, is indicated with a dashed border. Right: Visualisation of DAPNet (331 genes), with nodes coloured by direction of expression relative to albuminuria. Stacked bar plots show the proportion of DAPNet genes upregulated or downregulated with albuminuria (top), and the proportion of cis-eQTL, trans-eQTL, and non-eQTL genes within the network (bottom). (C) Scatter plots showing DAPNet eigengene correlation with scaled urinary albumin excretion (UAE) at weeks 8, 16, and 24. Spearman correlation coefficients and p-values are shown. (D) Bar plot showing the percentage of genes in each podocyte network that are albuminuria pGenes, with the ratio of pGenes to total network size annotated. DAPNet has the highest enrichment. Asterisks indicate significance (* p < 0.05, ** p < 0.01, *** p < 0.001).

Across the 12 podocyte networks identified, we found that 10 showed a significant correlation with albuminuria, and 7 correlated across all 3 timepoints (Figure 4B). One network stood out with 201 of 331 of its genes (60.7%) being downregulated in the setting of albuminuria (Figure 4C). This network showed the strongest negative correlation with albuminuria across the 12 networks and contained 12 albuminuria pGenes (Figure 4C and D). With these features, this network mirrors the signal observed in the albuminuria pQTL analysis, where most pGenes were downregulated in 129 mice (Figure 2E). We termed this network the Disease-Associated Podocyte Network (DAPNet, Extended Figure 5A) and found that DAPNet is enriched for vesicular transport, protein transport, and metabolic pathways such as cholesterol homeostasis and sphingolipid metabolism (Extended Figure 5B). Notably, DAPNet was the only podocyte network that showed significant enrichment for both microalbuminuria (adjusted p = 0.00149) and late-stage DN (adjusted p = 0.02061) in human GWAS. The leading genes driving this enrichment included Col4a3, Col4a4, and Snx6 for late-stage DN, and Plekho1, Tnfsf12, and Paqr7 for microalbuminuria, several of which showed consistent negative correlations with albuminuria across multiple timepoints (Extended Figure 5C).

Accordingly, we utilised DAPNet to prioritise a tractable number of candidate genes for functional proof-of-concept testing. Twelve of our albuminuria pGenes belong to DAPNet: Ankrd46, Dcaf6, Dcaf8, Ebag9, Ecd, Emc2, Igsf8, Ptprd, Rrm2b, Slc31a2, Tfb2m, and Znrf2. Their putative functions are shown in Suppl. Table 4. One of these, Ptprd, has been previously identified as a GWAS hit for T2D in Han Chinese and Mexican cohorts^37, 38^ and for hypertension^39,40^; the other genes have not been previously linked to diabetes or DN. As a control, we also selected Mllt3, a podocyte-expressed pGene linked to albuminuria and macrophage infiltration but not included in DAPNet, which has been previously associated with eGFR decline^41^ and hypertension^42^.

### Functional validation of DN-susceptibility genes

Albuminuria is a measurable consequence of impaired renal filtration capacity, and genes contributing to this trait are expected to regulate processes essential for podocyte function, including endocytosis, vesicle trafficking, and cytoskeletal dynamics. To directly assess the functional roles of 12 DAPNet candidate genes, we performed targeted knockdown in Drosophila nephrocytes, which are anatomically and functionally analogous to mammalian podocytes^43–45^. Only 6 of the DAPNet genes, Slc31a2, Ptprd, Emc2, Znrf2, Dcaf6, and Dcaf8 (along with Mllt3) had fly orthologs, were also eQTLs, and were therefore selected for functional validation and subjected to the assay. We used a Drosophila filtration assay based on the MHC-ANF-RFP reporter, in which muscle-secreted RFP is taken up by pericardial nephrocytes via endocytosis involving filtration via the nephrocyte slit diaphragm, accumulating as intracellular vesicles, providing a quantitative, visual readout of filtration capacity. Nephrocytes were labelled with Hand-GFP and the DotGal4 driver for cell-specific knockdown and visualisation (Figure 5A, C). To validate the sensitivity and dynamic range of the assay, we used Wingless (Wg), the Drosophila ortholog of Wnt1, part of a well-established pathway regulating cellular architecture in renal development and disease^46–51^. Knockdown of Wg resulted in a marked reduction in nephrocyte number and a near-complete loss of RFP uptake, confirming the assay’s sensitivity for detecting impaired filtration. As a negative control, we used the Hox gene Sex combs reduced (Scr^52^), which is not known to affect nephrocyte function. For each gene, RFP levels were normalised to Scr to allow quantitative comparison across conditions (Figure 5B).

**Figure 5.**
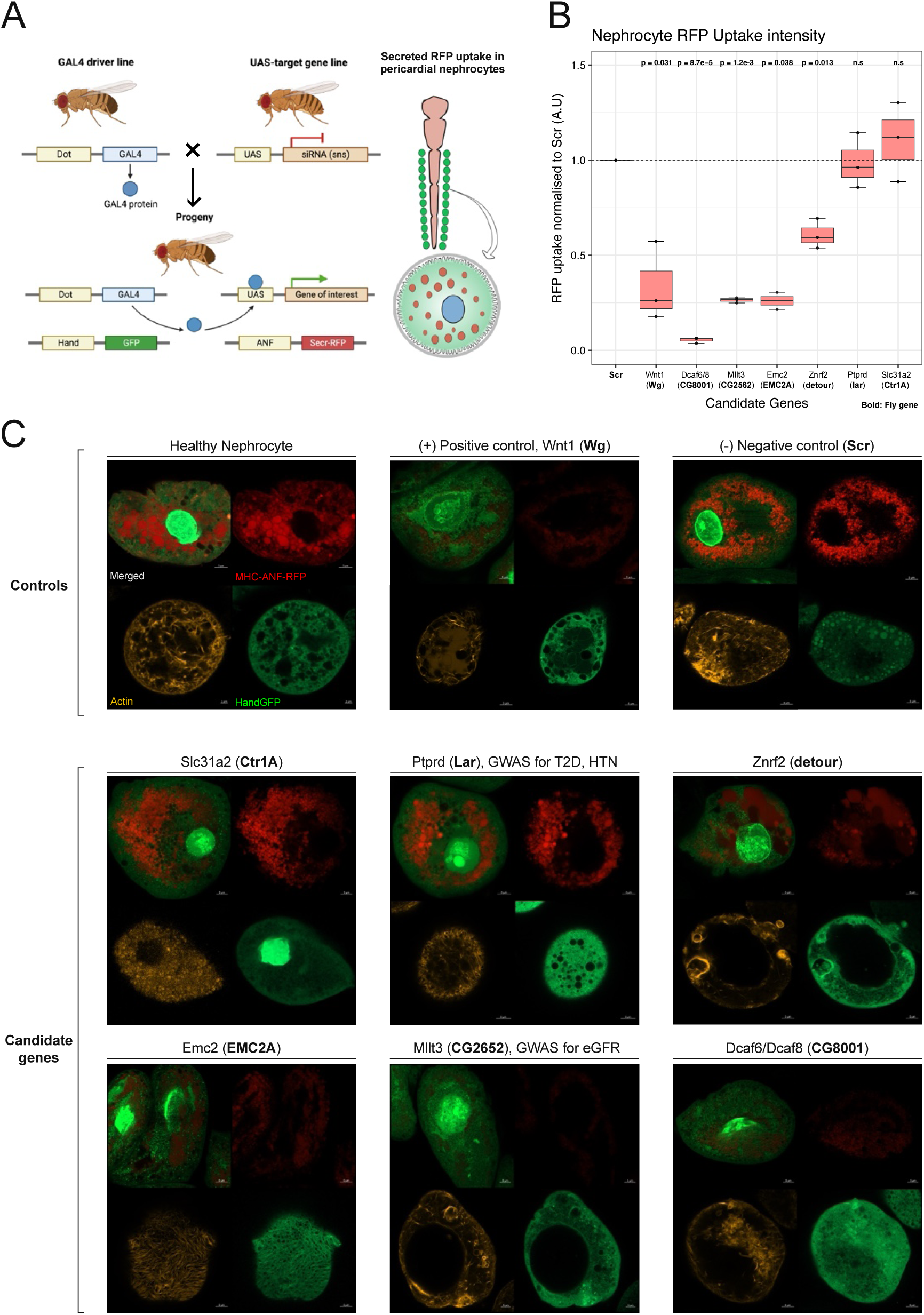
Knockdown of DAPNet genes disrupts nephrocyte filtration and actin organisation. (A) Representative schematic illustrating the nephrocyte assay system. (B) Boxplot showing RFP uptake by nephrocytes, normalised to the negative control gene (Scr), for each candidate gene knockdown condition. Significance was assessed using the Wilcoxon rank-sum test. (C) Representative confocal images of single nephrocytes from each knockdown condition, acquired at 63x magnification. Upper panels show MHC-ANF-RFP uptake; lower panels show filamentous actin distribution visualised using the FourColors reporter. Scale bars as indicated.

To assess gene function, DotGal4; Hand-GFP, MHC-ANF-RFP flies were crossed with UAS-RNAi lines targeting each candidate gene, and nephrocyte phenotypes were analysed in the resulting F1 progeny using confocal imaging. Gene knockdown resulted in distinct filtration phenotypes (Figure 5C). Knockdown of Mllt3, Znrf2, Dcaf6/8, and Emc2 produced significantly reduced filtration, whereas knockdown of Ptprd and Slc31a2 did not reduce overall filtration. However, in both Ptprd and Slc31a2 knockdown conditions, the number and size of endocytic vesicles were reduced, indicating altered vesicle dynamics. In contrast, Znrf2 knockdown resulted in loss of distinct vesicles and cytoplasmic leakage, suggesting severe disruption of cellular integrity. The most pronounced defects were observed in Dcaf6/8 and Mllt3 knockdown nephrocytes, which exhibited a near-complete loss of RFP uptake.

Endocytosis in nephrocytes is driven by actin filaments, which form the structural basis of slit diaphragms and regulate vesicular trafficking^53–55^. To assess whether candidate gene perturbation affected cytoskeletal organisation, we examined the distribution of filamentous actin in pericardial nephrocytes. In control cells, actin formed a dense and organised network surrounding lacunar channels, consistent with normal filtration architecture. In contrast, gene knockdown resulted in distinct cytoskeletal defects. Wg knockdown led to a fragmented actin network and enlarged peripheral lacunae. Dcaf6/8 knockdown resulted in a polarised and asymmetric distribution of actin. The most severe phenotypes were observed in Mllt3 and Znrf2 knockdown nephrocytes, where actin filaments were thinner, predominantly peripheral, and associated with the formation of large vacuole-like structures. Similar enlarged endocytic vesicles have been reported following Rab7 silencing^56^, suggesting disruption of endosomal trafficking pathways.

Notably, genes showing the most severe nephrocyte phenotypes upon knockdown were also among those with the strongest UACR associations in the human cohort. Mllt3, Dcaf6, and Znrf2, which produced pronounced filtration and cytoskeletal defects, each harboured multiple significantly associated variants in diabetic participants of the All of Us cohort (Extended Figure 6A, B, C), providing cross-species support for their role in podocyte dysfunction.

## Discussion

Inherited susceptibility to kidney injury in diabetes is complex, shaped by variation across multiple genes and cell types. Despite decades of human genetic studies, the causal genes regulating susceptibility to albuminuria, a powerful determinant of risk for progressive loss of kidney function in DN^57^, remain poorly defined. Early GWAS of albuminuria identified only CUBN as a robustly replicated genome-wide significant locus in general-population European-ancestry meta-analyses^15,58^. Subsequent studies were able to identify additional loci implicating genes involved in tubular transport, glomerular filtration barrier integrity, and TGF-β signalling, including SHROOM3, MUC1, COL4A3/A4, TGFB1, and NR3C2, among others^15,59^. Despite these advances, explained heritability of albuminuria and the translation of loci into biological mechanisms remain incomplete. Here, we present and validate a multi-layered systems genetics framework that integrates mouse QTL mapping, single-cell transcriptomics, gene coexpression network analysis, functional validation, and human genetic association to systematically prioritise candidate genes underlying albuminuria in DN. Across all analytical layers, a consistent signal implicating podocyte dysfunction has emerged. Specifically, our findings suggest that disruption of podocyte vesicle trafficking and actin cytoskeletal organisation may contribute to disease susceptibility. The identification of 26 novel pQTLs and 192 candidate genes (pGenes) for albuminuria allowed us to build a prioritisation pipeline for functional testing of novel candidates in DN. While this study focused on albuminuria as a primary trait, we provide a catalogue of pGenes/eGenes associated with macrophage infiltration and glomerulosclerosis, as well as 95 cell-type-specific coexpression networks across the full spectrum of kidney cell types.

Two of the 28 albuminuria loci identified were previously associated with albuminuria in independent mouse crosses, providing cross-population validation of our QTL approach. Furthermore, the enrichment of pGenes for human CKD and DN GWAS signals, including identification of 4 genes previously linked to DN in a Finnish population^33^, supports the relevance of our approach to human disease. The shared pQTL between macrophage infiltration and albuminuria on chromosome 4 points to common genetic regulatory mechanisms that may reflect the well-documented inflammatory contributions to DN pathogenesis^60^. Interestingly, one of the four significant pQTLs for urinary succinate levels mapped to a locus on chromosome 13 (13:74325867) containing Sdha, which encodes succinate dehydrogenase subunit A, the mitochondrial enzyme that catalyses the conversion of succinate to fumarate in the TCA cycle. The co-localisation of a succinate pQTL with the gene encoding the primary enzyme regulating succinate metabolism provides further confidence in gene-phenotype associations arising from QTL mapping. Overall, these observations validate the biological significance of our systems genetics approach for gene/pathway identification in DN.

The predominant expression of albuminuria pGenes in podocytes and their broad downregulation in the disease-susceptible 129 strain were striking and consistent findings. Podocyte loss and dysfunction are well-established early features of diabetic nephropathy^61–63^, and the broad transcriptional dysregulation we observed highlights the contribution of these genetic pathways in podocytes to DN-associated albuminuria. Furthermore, as contributors to DN pathogenesis, these pathways represent potential targets for therapy. This aligns with emerging evidence from human studies that podocyte-specific transcriptional programmes are disrupted early in diabetic kidney disease^64^.

Network analysis, combined with phenotypic information, has successfully identified disease-related modules, providing support for cellular pathways and underlying effector genes driving pathology^65–69^. We utilised network analysis here to construct cell-type-specific, disease-associated modules in DN. While we demonstrated that several podocyte coexpression modules show widespread correlation with albuminuria, only DAPNet passed the most stringent criterion of enrichment for human DN GWAS signals across both microalbuminuria and late-stage disease. This human GWAS enrichment distinguished DAPNet as most likely to reflect biologically conserved and clinically relevant disease processes. Accordingly, we used DAPNet to prioritise genes for functional testing.

The Drosophila nephrocyte has substantial genetic and structural similarities to the human podocyte and has become a well-accepted system for functional genetic testing in glomerular disease^43–45^. Of the 12 DAPNet albuminuria pGenes, 6 had fly orthologs (Slc31a2, Ptprd, Emc2, Znrf2, Dcaf6, and Dcaf8) and were selected for functional validation alongside Mllt3, a podocyte-expressed pGene associated with both albuminuria and macrophage infiltration that had been previously linked to eGFR decline and hypertension but is not a DAPNet member. With the exception of Ptprd, which has prior links to T2D and hypertension, the remaining candidates have no previously reported associations with albuminuria or kidney disease. Podocytes rely on regulated endocytosis and membrane trafficking to maintain the structural integrity of slit diaphragms and foot processes, and disruption of vesicle trafficking pathways has been directly implicated in proteinuric kidney disease across multiple models^70,71^. The enrichment of DAPNet for vesicular transport, protein trafficking, and cholesterol and sphingolipid metabolism is mechanistically coherent with this biology, as the podocyte slit diaphragm is assembled within cholesterol and sphingolipid-enriched lipid raft microdomains, and dysregulation of podocyte lipid metabolism causes cytoskeletal remodelling, endoplasmic reticulum stress, and apoptosis in experimental and human DN^61^.

Knockdown of each DAPNet gene caused abnormalities in nephrocyte structure or function, confirming their importance in podocyte biology and affirming the utility of network and pathway analysis for candidate prioritisation. Differences in phenotype severity reflected the distinct functional roles of each gene: Mllt3, Dcaf6/8, and Emc2 knockdown produced the most severe filtration deficits, with near-complete loss of RFP uptake, suggesting non-redundant roles in maintaining nephrocyte endocytic capacity; Znrf2 knockdown produced a qualitatively distinct phenotype characterised by cytoplasmic leakage and dissolution of vesicular structure, consistent with a role in maintaining membrane or endosomal integrity rather than vesicle formation; and Ptprd and Slc31a2 knockdown did not significantly reduce overall filtration, though vesicle dynamics were altered in both cases, suggesting these genes influence the efficiency rather than capacity of endocytosis. Together, these findings establish vesicle trafficking and actin cytoskeletal regulation as mechanistically central to the candidate gene set, validating the pathway enrichment identified in DAPNet.

The human genetic analysis in the All of Us Research Program provided a clinically relevant layer of validation of our systems genetics approach. 40% of eligible pGenes reached FDR-corrected significance for UACR association in 3,553 diabetic individuals. The convergence of MLLT3 and DCAF6 across fly functional validation and human FDR significance represents the strongest multi-species evidence for these genes as causal drivers of albuminuria susceptibility. MLLT3, previously associated with eGFR decline and hypertension^41,42^, emerges here as a podocyte-expressed gene whose loss impairs nephrocyte filtration and whose variants associate with UACR in diabetics, providing a mechanistic link between its known associations and glomerular function.

Among the DAPNet genes, DCAF6 and ZNRF2 are both E3 ubiquitin ligases, a class of proteins known to regulate and direct substrate proteins for degradation or functional modification^72,73^. While our demonstration of key roles for DCAF6 and ZNRF2 in podocytes is novel, ubiquitin ligases have been implicated in podocyte biology, regulating cytoskeletal dynamics, slit diaphragm integrity, and endosomal trafficking^74–79^. ZNRF2 has been linked to kidney injury^77^ and mTOR pathway activation^78^. Activation of the mTOR signalling pathway significantly impacts podocyte homeostasis and DN progression^80^. The severe filtration and cytoskeletal phenotypes observed upon knockdown of Znrf2 and Dcaf6/8 in Drosophila nephrocytes suggest that ubiquitin-mediated protein regulation plays a non-redundant role in maintaining podocyte homeostasis, providing a mechanistic hypothesis for why their downregulation in albuminuria-susceptible podocytes may contribute to glomerular dysfunction.

There are several limitations of this study. The Drosophila nephrocyte assay, while a powerful tool for testing cell-autonomous gene function, does not fully recapitulate the systemic diabetic and hypertensive context in which these genes act in the mammalian kidney. Genes that function through systemic or paracrine mechanisms may therefore be missed. The All of Us human genetic analysis was restricted to diabetic individuals with available UACR data, and the cohort, while diverse, was not specifically enriched for DN cases. Larger dedicated DN cohorts from different ethnic backgrounds would provide greater power to detect rare variant associations and to stratify by disease stage. Furthermore, the directionality of human variant effects relative to fly loss-of-function phenotypes could not be established without functional annotation of individual variants, limiting causal inference at the variant level in humans.

Beyond identifying specific candidate genes, this work has broader implications for understanding and managing CKD and DN. Our findings highlight pathways amenable to therapeutic intervention, particularly given the established pharmacological tractability of E3 ubiquitin ligases and their associated degradation machinery in other diseases. The accompanying catalogue of pGenes and cell-type-specific coexpression networks linked to macrophage infiltration and glomerulosclerosis extends the utility of this resource beyond albuminuria, providing a foundation for future work on the inflammatory and fibrotic components of DN. More broadly, our studies demonstrate the capability of cross-species systems genetics to uncover novel functional and genetic pathways in diabetic kidney disease. We have identified novel pQTLs for albuminuria, glomerulosclerosis, and macrophage infiltration, accomplished functional verification of novel candidate genes emerging from combined analysis of mouse and human datasets, and established a generalisable framework for converting genetic association signals into mechanistic insight, addressing a bottleneck that has long limited the clinical translation of human kidney disease genetics.

## Methods

### Animal Studies and Timepoint collections

F_2_ mice were produced by crossing the two parental lines, obtained as described previously^26,28^. All animal studies were reviewed and approved by the Institutional Animal Care and Use Committee of the National University of Singapore. Mice were housed in the Duke-NUS vivarium under controlled conditions, kept in a temperature-regulated environment with a 12-hour light/dark cycle, and had free access to water. The mice were fed a standard chow diet (Specialty Feeds SF100-100, irradiated rat and mouse cubes). F_2_ mice were raised for 24 weeks and placed in metabolic cages at 8, 16, and 24 weeks to collect urine over 24 hours. Albuminuria was measured using an enzyme-linked immunosorbent assay (Albuwell M, Exocell). Blood samples were obtained by puncturing the left lateral saphenous vein with a 25-gauge needle, collecting approximately 2-3 µl of blood^27^. Blood glucose levels were measured using a glucometer (Ascensia Bayer Elite). Mice were sacrificed at 24 weeks of age, and the kidneys were harvested for further analysis. One kidney was immediately frozen in liquid nitrogen, while the other was fixed in 10% neutral buffered formalin (NBF). The fixed kidneys were subsequently embedded in paraffin and sectioned at a thickness of 5 µm for histological examination.

### Single-cell RNA Sequencing on parental strains

Single-cell RNA sequencing was performed on whole kidney and glomerular cell isolates from 10-week-old age-matched wild-type (BL6WT and 129WT) and Akita-ReninTg (BL6AR and 129AR) mice. All mice were anaesthetised with intraperitoneal ketamine, perfused with chilled HBSS to flush blood from the kidneys, and subsequently with chilled Dynabeads. Kidneys were excised, decapsulated, and minced into 1 mm³ pieces.

#### Glomerular cell isolation

Minced tissue was digested in prewarmed enzyme solution 1 (1 mg/ml Collagenase I, 1X Pronase, and 0.33 µl/ml DNase in HBSS) for 10 min in a shaking 37°C water bath. The resulting suspension was filtered twice through 100 µm cell strainers, washed with chilled HBSS, and centrifuged at 300 × g for 4 min at 4°C. The pellet was resuspended in 1.5 ml HBSS, placed on a magnetic stand for 5 min, and the retained glomeruli-bead complexes were washed four times with chilled HBSS. To dissociate glomerular cells, the complexes were digested in prewarmed enzyme solution 2 (2% Trypsin, 2 U/ml Dispase, 2 U/ml Collagenase D, and 0.33 µl/ml DNase in HBSS) for 40 min at 37°C, with pipetting every 10 min and passage through a 27G needle at the 30 min mark. Digestion was halted with ice-cold 10% FBS in PBS and the suspension centrifuged at 300 × g for 4 min at 4°C. To separate beads from glomerular cells, the pellet was resuspended in ice-cold 3% FBS in PBS, placed on a magnetic stand for 3 min, and the supernatant collected. This step was repeated twice, with all washes pooled. The final suspension was centrifuged at 300 × g for 4 min at 4°C and resuspended in ice-cold 3% FBS in PBS. 15,000 glomerular cells per mouse were processed following the Chromium Single Cell 3’ Library and Gel Bead Kit v.3 protocol.

#### Whole kidney cell isolation

Minced tissue was digested in buffer containing 1 mg/ml Collagenase I and 0.33 µl/ml DNase in HBSS for 20 min at 37°C in a shaking water bath. The digested tissue was filtered through a 100 µm cell strainer, washed with chilled HBSS, and centrifuged at 300 × g for 4 min at 4°C. The pellet was resuspended in 2 ml chilled HBSS, mixed with 20 ml of 1X Red Blood Cell Lysis Solution, and centrifuged at 300 × g for 10 min at 4°C. The pellet was resuspended in HBSS and placed on a magnetic stand for 5 min to separate glomeruli, which were retained, from the tubular fraction, collected in the supernatant. The tubular fraction was digested in 2 ml digestion buffer for 15 min at 37°C, while concurrently the glomerular fraction was digested in 1 ml digestion buffer for 25 min, followed by magnetic separation to remove free beads. Both fractions were combined, centrifuged at 300 × g for 5 min at 4°C, and debris removed using Miltenyi Biotec’s Debris Removal Solution. The final pellet was resuspended in HBSS, filtered through a 30 µm cell strainer, centrifuged at 300 × g for 5 min at 4°C, and resuspended in 0.5% BSA in PBS. 20,000 whole kidney cells per mouse were processed according to the Chromium Single Cell 3’ Library and Gel Bead Kit v.3 protocol.

#### Preprocessing and quality control

Three batches of single-cell RNA sequencing were performed: two for whole kidney isolates and one for glomerular isolates. The two whole kidney batches were integrated for joint analysis. Whole kidney and glomerular datasets were first analysed and annotated separately, then combined for differential expression analysis and subsequent downstream work. Raw sequencing data were processed using the 10x Genomics Cell Ranger pipeline (v3.1.0) and aligned to the mm10 reference transcriptome. The resulting count matrices were imported into R and processed using the Seurat package (v4.0.3). Doublet removal was performed on each sample individually using scrubletR, the R implementation of the Python tool Scrublet, with default settings, and doublet annotations were retained in sample metadata for downstream filtering.

#### Preliminary analysis of whole kidney samples

Whole kidney samples were filtered to retain cells with 200–4,000 detected genes, fewer than 50% mitochondrial reads, fewer than 20% ribosomal reads, and fewer than 3% haemoglobin reads. Samples within each batch were merged into a single object, retaining only genes common to both batches and expressed across all samples. Cell cycle effects were accounted for by scoring each merged object using the CellCycleScoring function in Seurat, with scores subsequently regressed out during SCTransform normalisation. The two batch objects were then integrated following Seurat’s standard CCA workflow [282], using the SCT assay and 3,000 integration features. PCA was performed on the integrated object, and the first 50 principal components were used for clustering and UMAP visualisation. Clustering was performed at a resolution of 1 and UMAP generated using the RunUMAP function in Seurat. Cell types were manually annotated using established marker genes.

#### Preliminary analysis of glomerular samples

Glomerular samples were filtered to retain cells with 200–4,000 detected genes, fewer than 10% mitochondrial reads, fewer than 20% ribosomal reads, and fewer than 3% haemoglobin reads. Samples were merged into a single object and scored for cell cycle gene expression using the CellCycleScoring function in Seurat, with scores subsequently regressed out during SCTransform normalisation. PCA was performed on the merged object, and the first 50 principal components were used for clustering and UMAP visualisation. Clustering was performed at a resolution of 1 and UMAP generated using the RunUMAP function in Seurat. Cell types were manually annotated using established marker genes from the literature.

#### Combined analysis

For combined analysis, whole kidney and glomerular datasets were read into R as separate objects and filtered to remove previously identified doublets. Samples were merged batchwise to yield three objects, two for whole kidney and one for glomerular, and filtered according to the quality thresholds described above. Each object was scored for cell cycle gene expression using the CellCycleScoring function in Seurat, with scores regressed out during SCTransform normalisation. The three objects were then integrated using Seurat’s Reciprocal Principal Component Analysis (RPCA) method on the SCT assay. Cell type identities were assigned based on annotations derived from the preliminary single-extract analyses.

### Urine Organic Acid Metabolomics

Urine samples were processed for targeted organic acid quantification by gas chromatography-mass spectrometry (GC-MS). Briefly, 300 µL of tissue homogenate or cell suspension was combined with 10 µL of internal standard and 10 µL of O-ethylhydroxylamine hydrochloride (Sigma Aldrich, USA) to protect alpha-keto groups prior to extraction. Organic acids were extracted into 500 µL of ethyl acetate and subsequently derivatised with N,O-Bis(trimethylsilyl)trifluoroacetamide to form trimethylsilyl derivatives. Separation was performed by gas chromatography on an Agilent Technologies HP 7890A instrument fitted with a VF-1ms capillary column (15 m × 250 µm × 1 µm) at a constant flow rate of 2.5 ml/min at 23.3 psi. The GC temperature programme was as follows: initial hold at 70°C for 2 minutes, ramp at 40°C/min to 300°C, and final hold at 300°C for 2.25 minutes. Quantification was performed by selected-ion monitoring on an Agilent 5975C mass spectrometer using stable-isotope dilution. Metabolite concentrations were normalised to urinary creatinine levels to account for variation in urine concentration across samples.

### Histology and immunohistochemistry

#### Periodic Acid-Schiff staining

FFPE kidney sections from F_2_ mice were used for Periodic Acid-Schiff (PAS) staining to assess kidney damage. PAS staining detects glycogen, glycoproteins, and mucosal polysaccharides, allowing visualization of glomerular scarring and tubular atrophy characteristic of diabetic nephropathy. Staining was performed following the manufacturer’s instructions (Abcam, PAS Staining Kit, ab150680). Briefly, sections were deparaffinized in xylene, rehydrated through graded alcohols, treated with Periodic Acid solution for 10 minutes, rinsed in distilled water, and incubated in Schiff’s reagent for 30 minutes. After washing in hot running tap water, sections were counterstained with Hematoxylin, dehydrated, and mounted with D.P.X solution.

#### CD68+ Immunohistochemistry

FFPE kidney sections from F_2_ mice were also used to quantify macrophage infiltration via CD68 staining. Sections were deparaffinized, rehydrated, and subjected to antigen retrieval by pressure-treating in a pH 9 Tris-EDTA buffer for 45 minutes at 50 kPa. After PBS washing, sections were incubated with BLOXALL solution (Vector Laboratories, SP-600) for 10 minutes to block endogenous peroxidase activity, followed by another PBS wash. Non-specific binding was blocked with 2.5% horse serum for 20 minutes before incubation with CD68 antibody (1:100, Cell Signaling, 97778). Following PBS washes, sections were incubated with IMMPress HRP horse anti-rabbit IgG polymer (Vector Laboratories, MP-7401) for 30 minutes. Sections were incubated with the Pierce DAB substrate (ThermoFisher, 34002) for 20 minutes, or until any signal was observed. Finally, sections were counterstained with Haematoxylin, dehydrated, and mounted with D.P.X solution.

### Histological Examination

Kidney histopathology was evaluated by pathologists blinded to group allocation using a semi-quantitative scoring system adapted from the criteria for human diabetic nephropathy (DN)^31^. Mesangial expansion, nodular sclerosis, and global glomerulosclerosis were scored, and individual feature scores were summed to obtain a total score (Suppl. Table 1). Mesangial expansion was graded by the number of mesangial cells in areas of expansion: <4 = 0, 4–5 = 1, 6–7 = 2, 8–10 = 3, ≥11 = 4. Nodular sclerosis and thickening of the tubular basement membrane in non-atrophic tubules were scored as absent (0) or present (1). Global glomerulosclerosis was graded as absent (0), ≤50% (1), or >50% (2). Interstitial fibrosis and tubular atrophy were scored by cortical involvement: absent (0), <25% (1), 25–50% (2), >50% (3). Arteriolar hyalinosis was scored as absent (0), one arteriole (1), or >1 arteriole (2).

### Identification of extreme phenotypes

The F_2_ mice represent a diverse group of samples. To stratify the samples, we used the distribution of albuminuria to identify those at the extremes. This was done to find mice with consistently high or low albuminuria across all three time points. Urinary albumin levels were first log-transformed and z-scaled at each time point, converting the values into z-scores. We then grouped samples that consistently had z-scores above 1 or below -1 (which corresponds to 1 standard deviation above or below the mean) across all three time points. This resulted in a total of 26 mice, with 13 at each extreme, hereafter referred to as the “High Albuminuria” and “Low Albuminuria” groups.

### Whole genome sequencing and variant calling

Tail DNA was collected at 12 weeks of age and used for whole-genome sequencing (WGS). DNA was extracted using the Omega Bio-Tek E.Z.N.A Tissue DNA kit (D3396-00S), following the manufacturer’s instructions. Sequencing aimed at a minimum depth of 13x. The resulting FASTQ files were processed according to the Genome Analysis Toolkit (GATK) v4 best practices pipeline for germline short variant calling of SNPs and indels. Briefly, FASTQ files were aligned to the Mus musculus GRCm38 primary assembly reference genome (Ensembl Release 102) using the BWA-MEM algorithm. BAM files were sorted and indexed with samtools v1.16.1, then duplicates were removed using the MarkDuplicates. BAM files were re-sorted and indexed again with samtools to improve efficiency in subsequent steps.

Variant calling was performed using three reference variant sets: (i) the combined Mouse Genome Project (MGP) variant set, (ii) the C57BL/6NJ variant set, and (iii) the 129SvEvBrd variant set, sourced from the <u>dbSNP 142 release</u>. Base recalibration was conducted in two steps. First, BaseRecalibrator was run to build a model for each sample, accounting for factors such as the read group, sequencer-assigned quality scores, sequencing cycle position, and sequence context. Next, ApplyBSQR was used to recalibrate the base scores based on this model, assigning each read a new quality score. This step is crucial for improving variant calling accuracy and reducing errors introduced by sequencing artefacts. Following recalibration, we ran HaplotypeCaller to call variants for each sample. This tool calls SNPs and indels simultaneously using local de novo assembly, generating a Genotype Variant Calling File (GVCF) for each sample. These GVCF files were combined into a GenomicsDB database, which efficiently stores and queries large genomic datasets. Joint genotyping was then performed across all samples using GenotypeGVCFs, producing a single VCF file containing the genotypes for all samples. The merged VCF file was further refined using VariantRecalibrator to improve call accuracy, producing a tranches file. Variants were filtered using this tranches file to remove low-quality calls. The final VCF file was annotated using the Ensembl Variant Effect Predictor (VEP) with [Ensembl Release 100](https://ftp.ensembl.org/pub/release-100/variation/vcf/mus_musculus/) to provide functional annotations, including gene names, predicted protein effects, and conservation scores.

### Quantitative Trait Loci (QTL) mapping

#### Preprocessing

VCF files were filtered to remove parental strains and low-quality variants. The VCF file was then converted to a PLINK format using the command-line plink v1.9 tool. The VCF file was filtered to retain biallelic SNPs and exclude variants with a minor allele frequency (MAF) less than 0.05, a missing genotype rate greater than 0.1, and a Hardy-Weinberg equilibrium p-value less than 0.001. This resulted in 4,867,965 biallelic SNPs for downstream analysis.

#### Block construction and pruning

Haplotype blocks were constructed using the Haploview^81,82^ interpretation to identify blocks of SNPs in strong linkage disequilibrium (LD) with each other. This was done using the plink --blocks command, run separately for each chromosome, with the maximum size set to 130 Mb (corresponding to the length of the largest mouse chromosome, Chr1). SNPs were considered to be in LD if the bottom 90% confidence interval was greater than 0.7, and the top of the interval was at least 0.98. LD blocks were constructed using the indep-pairphase modifier in the plink command-line software, with a window size of 15 megabases (Mb), a step size of 1 base, and an R2 threshold of 0.8. The pairphase modifier uses values based on maximum likelihood phasing pruning SNPs to the same blocks as those constructed earlier. LD pruning reduced the number of SNPs from 4,867,965 to 240,507.

### Phenotype association mapping

Quantitative trait loci mapping of phenotypes was primarily done using GEMMA (Genome-wide efficient mixed model analysis)^83,84^. LD-pruned SNPs were used to construct the relatedness matrix from PLINK-formatted files. This matrix was then passed to the GEMMA command-line tool to perform association testing using Wald’s test.

### CD68+ quantification and QTL mapping

All F2 mice were assessed for CD68+ macrophage infiltration in the kidney. Kidney sections were stained as previously described, and images were captured using a 10x objective on a Nikon Eclipse Ni-E microscope. Three images of the cortex region were taken for each kidney section. The CD68+ area was measured from kidney cortex images using a custom macro in Fiji (ImageJ). Images were first processed with the H DAB colour deconvolution vector to isolate the DAB signal corresponding to CD68+ staining. Background was removed from the deconvolved DAB channel using a rolling ball radius of 20 pixels. To normalise staining by tissue area, two binary masks were generated per image. The first applied a fixed threshold (0–254) to determine tissue area. The second used a dynamic threshold identified by locating the first histogram bin above an intensity of 95 containing more than 1,000 pixels, capturing the CD68+ signal. Both masks were converted into binary images and quantified using Fiji’s Measure function. The CD68+ area was normalised to the tissue area and averaged across three images per sample. The normalised CD68+ area was used as a phenotype for QTL mapping, employing GEMMA as described earlier.

### Association mapping for the proportion of sclerosed glomeruli

QTL mapping of glomerulosclerosis was conducted using the proportion of sclerosed glomeruli as a quantitative trait. This proportion was calculated by dividing the number of scarred and damaged glomeruli by the total number of glomeruli examined. Because the data distribution was not normal and exhibited a zero-inflated property, we chose to use zero-inflated gamma regression to better model the data and perform association testing with the R package glmmTMB^85,86^. For each SNP, a full model including the SNP dosage was compared to a null model without the SNP term, using a likelihood ratio test (LRT), while incorporating the first five principal components (PCs) as covariates. Markers lacking genotype variation were excluded from this analysis. The likelihood ratio statistic was calculated as:

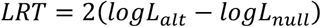

We derived genome-wide empirical significance thresholds using permutation testing. Briefly, the sclerosis data were randomly assigned to samples while keeping the genotype matrix and covariates unchanged, and the genomic association was performed. For each permutation, the maximum LRT statistic across all SNPs was recorded, with a total of 10,000 permutations to ensure stability. Empirical p-values were then calculated as:

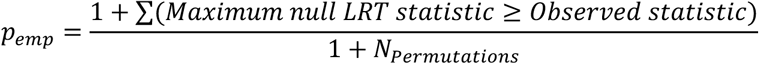

SNPs were assigned empirical p-values based on this null distribution and tested for significance (empirical p < 0.05).

### Hit prioritisation and candidate gene identification

Wald test p-values from GEMMA were adjusted for multiple testing using the Benjamini-Hochberg (BH) method. SNPs with adjusted p-values less than 0.05 were considered significant QTLs associated with albuminuria. Since the analysis was conducted on linkage disequilibrium (LD)-pruned SNPs, each significant marker represents a genomic region likely linked to the phenotype. To refine these regions, haplotype blocks defined by D’ were used to expand each significant SNP into its corresponding block. These blocks often spanned multiple megabases and included numerous genes, although only a subset was expected to be causal. To prioritise candidate genes, parental genotype data were used to identify loci that differed between the 129AR and BL6AR strains, focusing on genes with strain-specific sequence variation that may contribute to diabetic nephropathy susceptibility. Genes located within significant haplotype blocks were filtered to retain only those with sequence differences between 129AR and BL6AR mice. This list was further refined to include only genes with human orthologues and those expressed in the mouse kidney.

### RNA extraction and sequencing

RNA was extracted using the Omega Bio-Tek E.Z.N.A MicroElute Total RNA kit (R6831-02). Fresh frozen kidneys were cut on ice into 5 mg pieces, homogenised by adding 300μL of lysis buffer (from the extraction kit) and 1mm glass beads (ThermoFisher, 465945000), then pulsed in a bead beater for 60 seconds at 3,000 RPM. Tubes were inspected to ensure complete homogenisation, and if necessary, pulsed again. The supernatant was collected and processed according to the manufacturer’s protocol. The RNA was eluted in 20μL of water and stored at -80^∘^C. RNA integrity and sample purity were assessed using the Agilent Bioanalyzer 2100, and only samples with an RNA integrity number (RIN) greater than 7 were selected for sequencing. RNA sequencing was performed on an Illumina NovaSeq X platform in a 150 bp paired-end configuration.

### Sequence alignment and preprocessing

FASTQ files were initially trimmed using the Trim Galore tool to remove low-quality reads and adapter sequences^87^. The trimmed reads were then aligned to the Mus musculus GRCm38 primary assembly reference genome using the STAR aligner^88^. The Ensembl Release 100 gene GTF files were used for annotation, and a count matrix was generated with FeatureCounts^89^. The count matrix was imported into R and processed using the DESeq2 package. The data were filtered to retain genes with at least 10 counts in a minimum of 25% of the samples.

### Differential expression analysis

Differential expression analysis was performed using the DESeq2 package^90^. We aimed to identify genes that vary with albuminuria and employed two approaches. First, we used the extreme stratification groups to find genes differentially expressed between the high and low albuminuria groups. The DESeq function and the Wald test were used to detect gene expression changes. The second approach involved analysing continuous albuminuria levels across all samples. Here, the DESeq function and the likelihood ratio test were employed to find genes differentially expressed with albuminuria. Differential expression analysis was conducted separately for each timepoint, and the results were combined to identify a core set of genes associated with albuminuria. Wald test p-values were adjusted using the Benjamini-Hochberg method to account for multiple testing. Genes with adjusted p-values ≤ 0.05 were considered differentially expressed genes (DEGs). Pathway enrichment analysis was carried out using enrichR^91^.

### Expression Quantitative Trait Loci mapping

Gene expression traits were derived from variance-stabilized transformed (VST) RNA-seq counts, retaining genes with at least 10 counts in 25% of the samples. Genome-wide association testing between pruned SNP genotypes and gene expression traits was performed using GEMMA as described previously. Association testing was conducted under an additive genetic model using the LMM option (-lmm 4). Nominal p-values were derived from Wald tests of the SNP fixed effect. For computational efficiency, only SNP–gene pairs with nominal p < 1 × 10-2 were retained in the exported association results used for downstream processing. For each SNP–gene pair, genomic distance was calculated as the absolute difference between SNP position and the midpoint of the corresponding gene. Cis-eQTLs were defined as variant–gene pairs located on the same chromosome within ±1 Mb of the gene midpoint. All other associations were classified as trans-eQTLs. Cis-eQTL significance was assessed at the gene level. For each gene, the minimum nominal p-value among all cis SNPs was identified. Multiple testing correction was performed across all genes using the Benjamini–Hochberg false discovery rate (FDR) procedure. Genes with FDR < 0.05 were declared significant cis eGenes. For reporting purposes, the lead SNP (smallest nominal p-value) per significant cis eGene was retained. Trans-eQTL significance was assessed at the SNP–gene pair level. To control the family-wise error rate across trans associations, the total number of trans hypothesis tests was estimated as:

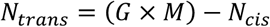

where G is the total number of genes tested, M is the total number of SNPs included in the GEMMA analysis, and *N*_cis_ represents the number of variant–gene pairs located within ±1 Mb of gene midpoints. The number of cis pairs was calculated by genomic interval overlap between SNP coordinates and gene-centred ±1 Mb intervals. A Bonferroni-corrected significance threshold was defined as:

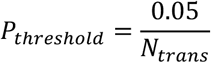

Trans SNP–gene pairs with nominal p-values below this threshold were declared significant. This correction reflects the full trans search space and is independent of nominal p-value filtering applied to the exported association results.

### Cell type-specific network construction

Gene coexpression networks were constructed using WGCNA, an approach that builds networks based on correlations in gene expression levels across samples^92^. Single-cell data from the 129AR mouse were used to identify differentially expressed genes (DEGs) for each cell type compared to all other cell types, filtered for log-fold change ≥ 1 and an adjusted p-value ≤ 0.05. This produced DEGs for each cell type, which were then used to filter the bulk RNA matrix containing gene expression data from the F2 mice. This method enabled us to construct gene coexpression networks with known single-cell expression patterns and identify genes co-expressed within the same cell type. For each cell type, the following steps were performed: first, the list of DEGs was used to filter the matrix, which was then normalised using DESeq’s variance-stabilising transformation. The normalised and filtered matrix was used to compute a soft-thresholding power using the pickSoftThreshold function. The soft thresholding power is a parameter that controls the degree of connectivity among genes in the network. A higher value results in a more connected network, while a lower value results in a sparser network. The pickSoftThreshold function automatically determines the optimal value.

The adjacency matrix was then computed using the adjacency function with the chosen value. This matrix was transformed into a topological overlap matrix (TOM) using the TOMsimilarity function, which measures the similarity between genes based on their connectivity within the network. Hierarchical clustering was performed on the TOM matrix using the hclust function with the ward.D2 method. The resulting dendrogram was segmented into modules with the cutreeDynamic function, which identifies modules of co-expressed genes based on their connectivity. The dendrogram was cut at a height of 0.25 (corresponding to a correlation of 0.75), merging similar modules. A minimum module size of 30 genes was set to define the modules. The module eigengene (ME) was calculated for each module using the moduleEigengenes function, representing the first principal component of the gene expression data within that module. The resulting modules, module eigengenes, and gene lists were named according to their cell type and the relative size of the network, with 1 indicating the largest, 2 the second largest, and so on.

### Network characterisation and MAGMA enrichment

Networks were annotated using several complementary approaches. First, network eigengenes were correlated with albuminuria at each time point using the cor function in R, with Benjamini-Hochberg correction applied within each cell type. We also quantified, for each network, the number of genes that were (i) albuminuria pGenes, (ii) differentially expressed between the 129 and BL6 strains, and (iii) eGenes identified from the eQTL analysis. To assess clinical relevance, networks were further tested for enrichment of human diabetic nephropathy GWAS signals using MAGMA (Multi-marker Analysis of GenoMic Annotation), which aggregates SNP-level summary statistics into gene-level associations while accounting for linkage disequilibrium. Because no human GWAS focused specifically on albuminuria was available, we used a published GWAS of diabetic nephropathy in which patients were stratified by albuminuria status. This dataset included comparisons between healthy controls and diabetic nephropathy cases across multiple stages, including microalbuminuria, macroalbuminuria, and end-stage renal disease. Summary statistics were obtained from the JDRF Diabetic Nephropathy Collaborative Research Initiative GWAS resource. GRCh37 gene location files and European reference data from the 1000 Genomes Project were obtained from the MAGMA reference portal. MAGMA analysis was performed in three steps. First, SNPs were annotated to genes using a 10-kb upstream and downstream window. Second, gene-level association statistics were calculated using the GWAS summary statistics together with the reference genotype data and SNP-to-gene annotation files. Third, gene set enrichment analysis was performed using the genes within each network. Networks with adjusted p-values below 0.05 were considered significantly enriched for the corresponding human trait.

### Drosophila Nephrocyte Imaging Assay

#### Generation of Dotgal4, UAS-FourColors line

The lines for Nephrocyte expression studies: dorothy-GAL4 (dotGAL4), myosin heavy chain (MHC) enhancer driving a red fluorescent protein (RFP) reporter gene fused with the Atrial Natriuretic Factor (ANF) secretion peptide or MHC-ANF-RFP, and hand enhancer driving green fluorescent protein (hand-GFP) were kind gifts from Zhe Han^93^. FourColors was generated by DNA synthesis (Synbio Technologies, NJ, USA) containing fluorescent protein sequences flanked by attL1/2 sites for Gateway recombination technology cloning (Invitrogen, USA) into the Drosophila expression vector pUASg.attB.3XHA (A kind gift from J. Bischof and K. Basler, Zurich^94^. The sequence was designed to express SV40 Nuclear Localisation Sequence fused to TagBFP^95,96^, Src Myristylation Sequence fused to mNeonGreen^97,98^, LifeActin fused to mKO^99,100^ and mKate2^101^ fused to the fly jupiter gene coding sequence for localization to microtubules^102^. The polycistronic construct contained P2A and T2A sequences separating each coding gene to allow for self-cleaving peptides^103^ as well as FRT sequences for use in Flp-dependent recombination^104^. The construct was injected into attP2 (BDSC Strain 8622) P[CaryP]attP2 68A4 by Best-Gene Inc. (California). Positive lines were crossed by standard genetics to dotGAL4 lines to generate stable combination lines expressing all four fluorescent proteins in nephrocytes.

#### Fly genetics and crosses

All stocks were obtained from the Bloomington Drosophila Stock Center (NIH P40OD018537). The TRiP lines used are summarised in Suppl. Table 17^105^.

#### Tracer uptake functional assay and Actin distribution in Drosophila nephrocytes

Female flies from F_1_ generation of fly cross Dotgal4; HandGFP; MHC-ANF-RFP x UAS RNAi line or Dotgal4; Four colors x UAS RNAi line were collected for dissection for tracer uptake experiments and actin distribution studies, respectively. Under anesthesia, fly appendages were removed and then transferred to a dissecting dish containing 1x PBS. The flies were placed ventral side up; the thorax and posterior end were removed using Dumont #5 forceps. With the abdomen open at both ends, a straight cut was made from anterior to posterior opening across the abdominal centerline, thereby unfolding the abdomen and removing the inner organs. The heart tube and pericardial nephrocytes, still attached to the cuticle, were immediately mounted for live imaging. The sample was placed cuticle-side down on a microscope slide with two layers of Scotch tape on either side. Glass coverslips were placed on the sample with a few drops of PBS, and the edges were sealed before imaging. Nephrocytes were imaged using a Zeiss LSM 800 confocal microscope with 40x and 63x objectives to visualise the nephrocyte arrangement, and in Airyscan mode for single-nephrocyte super-resolution imaging.

### Human variant level association analysis

Albuminuria candidate genes (pGenes) were tested for variant-level association with Urinary Albumin to creatinine ratio (UACR) in diabetic participants from the All of Us Research Program Controlled Tier Dataset v8. Participants were included if they had a recorded diagnosis of type 2 diabetes (SNOMED 201820, 201826). Participants prescribed metformin were excluded as metformin has been shown to independently reduce albuminuria through AMPK-mediated pathways^106^, which could confound the detection of genotype-level effects on UACR. Participants prescribed SGLT2 inhibitors were similarly excluded, as these medications directly reduce urinary albumin excretion independently of underlying glomerular pathology, potentially masking true genetic associations^107^. Participants with missing covariates (age, sex, race, or blood glucose) were further excluded, leaving 3,553 participants for analysis. Variants within the pGenes were obtained from gnomAD, and carrier status was queried from the All of Us database.

For each gene, covariate-adjusted linear regression was performed for all variants carried by at least 10 individuals in the diabetic cohort, with log₁₀-transformed UACR as the outcome and age, sex, race, and blood glucose as covariates. Within-gene multiple testing was controlled using the Benjamini-Hochberg procedure. Variants with adjusted p < 0.05 were classified as adverse (β > 0, higher UACR in carriers) or protective (β < 0, lower UACR in carriers), and the directional balance per gene was defined as the number of adverse minus protective variants. To assess whether the observed directional enrichment exceeded chance, a permutation procedure was implemented. Covariate-adjusted residuals of log₁₀-UACR were permuted 500 times using gene-invariant random seeds, ensuring the same phenotype shuffle was applied across all genes within each replicate. Per-variant statistics were computed using a closed-form ordinary least squares estimator on the residualised outcome, with within-gene Benjamini-Hochberg correction applied identically in each replicate. The empirical p-value per gene was defined as the proportion of permutations in which the number of nominally significant variants was equal to or exceeded the observed count, computed as (k + 1)/(N + 1). Empirical FDR was obtained by applying the Benjamini-Hochberg procedure across all 179 human orthologous genes. Genes were classified as empirical FDR < 0.05 (n = 72), empirical p < 0.05 (n = 55), or non-significant (n = 52).

## Author Contributions

T.M.C., E.P. and J.B. conceived the project. J.B. and T.M.C co-supervised the project and obtained funding. K.M. performed all bioinformatics analyses, including QTL and eQTL mapping, single-cell transcriptomic analysis, gene coexpression network construction, and human variant association analysis; carried out CD68 immunohistochemistry staining and quantification, and wrote the manuscript with help from J.B. and T.M.C. R.B.S. performed mouse breeding and F2 cross generation, coordinated sample collection, prepared single-cell suspensions for sequencing, and processed urine samples for metabolomics. B.L.F. scored and annotated the kidney histology sections. M.S.T. collected mouse phenotype data. J.W. prepared kidney tissue and performed RNA extraction for bulk RNA sequencing. S.S. performed Drosophila nephrocyte RNAi crosses and functional assays. J.C. and J.-P.K. performed GC-MS acquisition of urinary organic acids and contributed to metabolite interpretation. J.G. contributed to early-stage bioinformatics analysis and provided input. K.M., R.B.S., S.S., J.W., B.L.F., J.G., J.C., M.S.T., J-P.K., S.B.G., E.P., N.S.T., T.M.C., and J.B. contributed to data interpretation. All authors have seen and approved the final version of the manuscript.

## Acknowledgements

We thank the members of the Behmoaras lab for helpful discussions and insights. We are grateful to the Duke-NUS Vivarium staff for animal husbandry support. We acknowledge the All of Us Research Program participants and investigators for making data available for this study. The All of Us Research Program is supported by the National Institutes of Health Office of the Director. We thank Z. Han for providing the MHC-ANF-RFP, Hand-GFP, and DotGal4 fly lines, and J. Bischof and K. Basler for the pUASg.attB.3XHA expression vector. Fly stocks were obtained from the Bloomington Drosophila Stock Centre (NIH P40OD018537).

## Extended figures

**Extended Figure 1.**
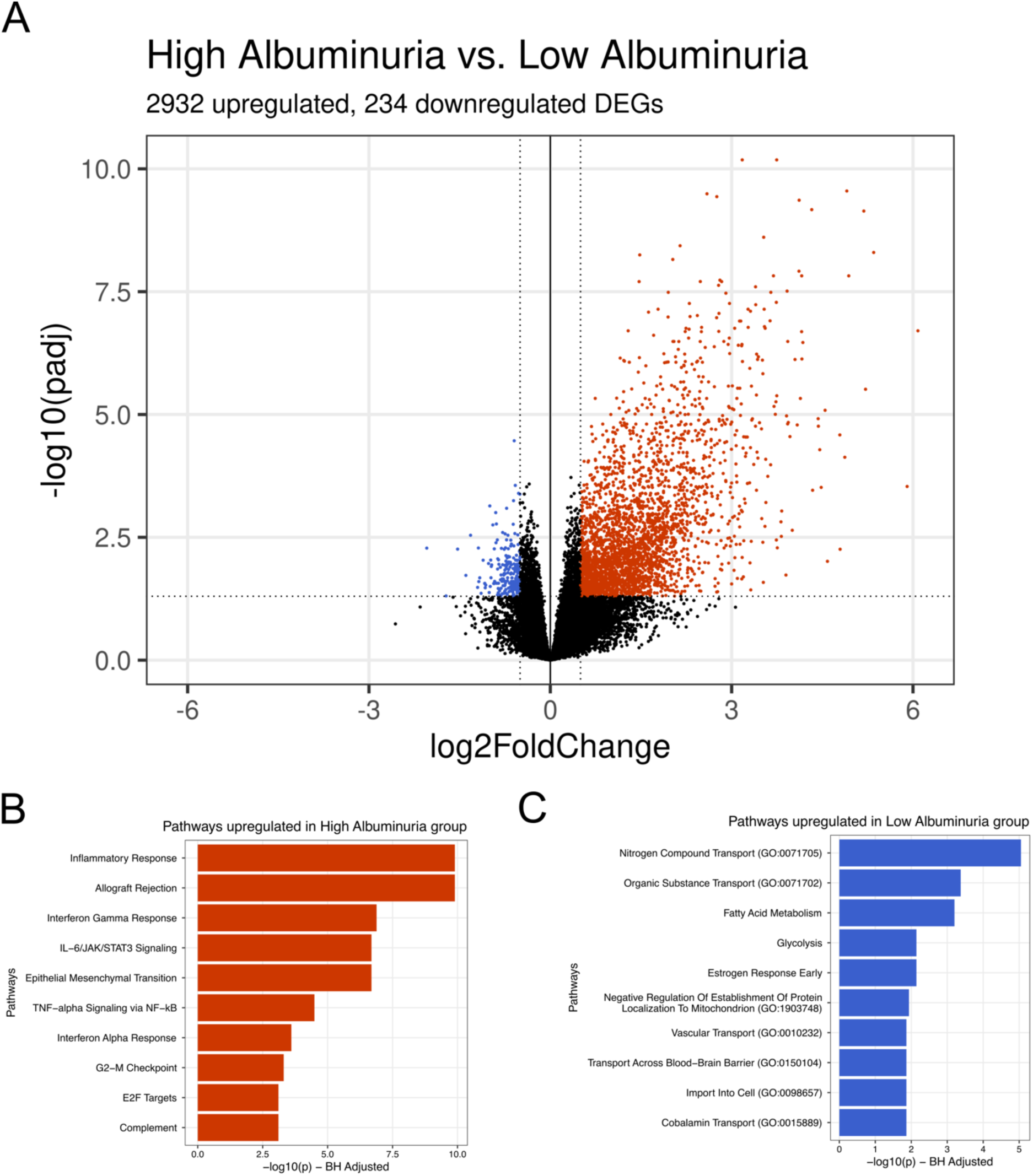
Differentially expressed genes (DEGs) between high and low albuminuria groups. (A) Volcano plot displaying the number of upregulated and downregulated DEGs between the two conditions. Significant genes (p.adj < 0.05, logFC >= 1) are colored in red (for upregulated genes in high albuminuria) and blue (for downregulated genes). (B) and (C) show pathways enriched in the upregulated and downregulated genes. Pathway enrichment was performed using enrichR.

**Extended Table 1.** eQTL mapping results by chromosome in the F2 kidney. For each chromosome, the number of significant cis-eGenes (FDR < 0.05) and trans-eGenes (Bonferroni-corrected) identified by linear mixed model eQTL mapping in whole kidney RNA-seq data from F2 mice (n=279) is shown. The final column indicates the number and percentage of cis-eGenes on each chromosome that are also albuminuria pGenes, defined as genes within significant pQTL loci after prioritization filtering. Percentages are calculated relative to the total number of cis-eGenes per chromosome. No albuminuria pGenes overlapped with trans-eGenes on any chromosome.

| Chromosome | cis-eGenes | trans-eGenes | pGenes & cis-eGenes |
| --- | --- | --- | --- |
| 1 | 358 | 5 | 28 (7.8%) |
| 2 | 387 | 1 | 0 |
| 3 | 279 | 0 | 1 (0.4%) |
| 4 | 291 | 4 | 14 (4.8%) |
| 5 | 367 | 1 | 0 |
| 6 | 325 | 4 | 7 (2.2%) |
| 7 | 578 | 9 | 0 |
| 8 | 310 | 1 | 1 (0.3%) |
| 9 | 307 | 2 | 0 |
| 10 | 221 | 5 | 1 (0.5%) |
| 11 | 443 | 7 | 0 |
| 12 | 227 | 1 | 0 |
| 13 | 257 | 2 | 0 |
| 14 | 169 | 0 | 12 (7.1%) |
| 15 | 187 | 4 | 16 (8.6%) |
| 16 | 172 | 1 | 0 |
| 17 | 313 | 1 | 2 (0.6%) |
| 18 | 140 | 1 | 0 |
| 19 | 163 | 1 | 0 |
| **Total** | **5494** | **50** | **82 (1.5%)** |

**Extended Figure 2.**
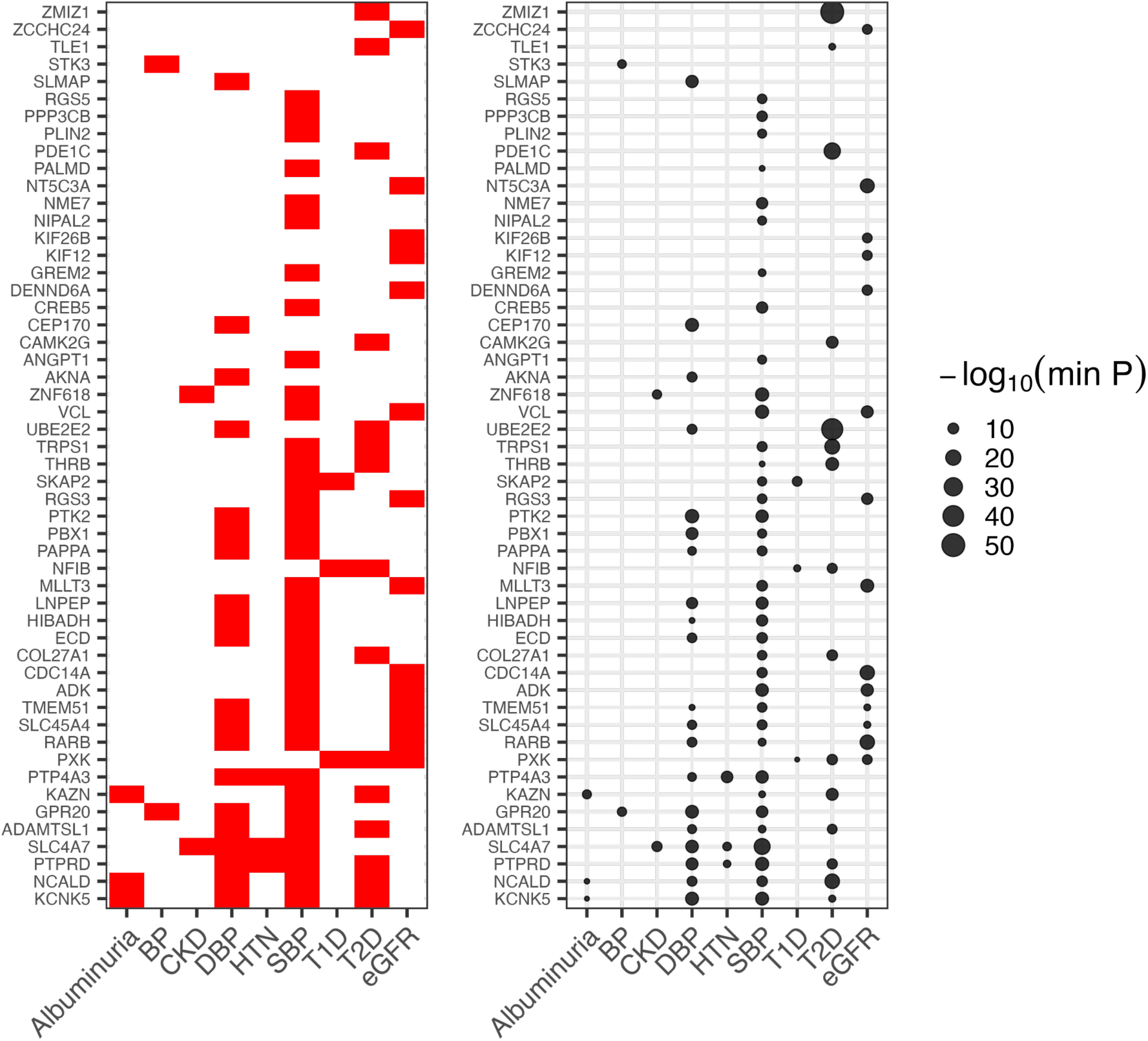
Albuminuria pGenes that are GWAS hits. Human orthologs of the genes have been plotted for convenience. Left: Plot showing which genes are GWAS hits across one or more categories. Right: Dotplot showing the lowest p-value for the SNP within the gene that is associated with the trait.

**Extended Figure 3.**
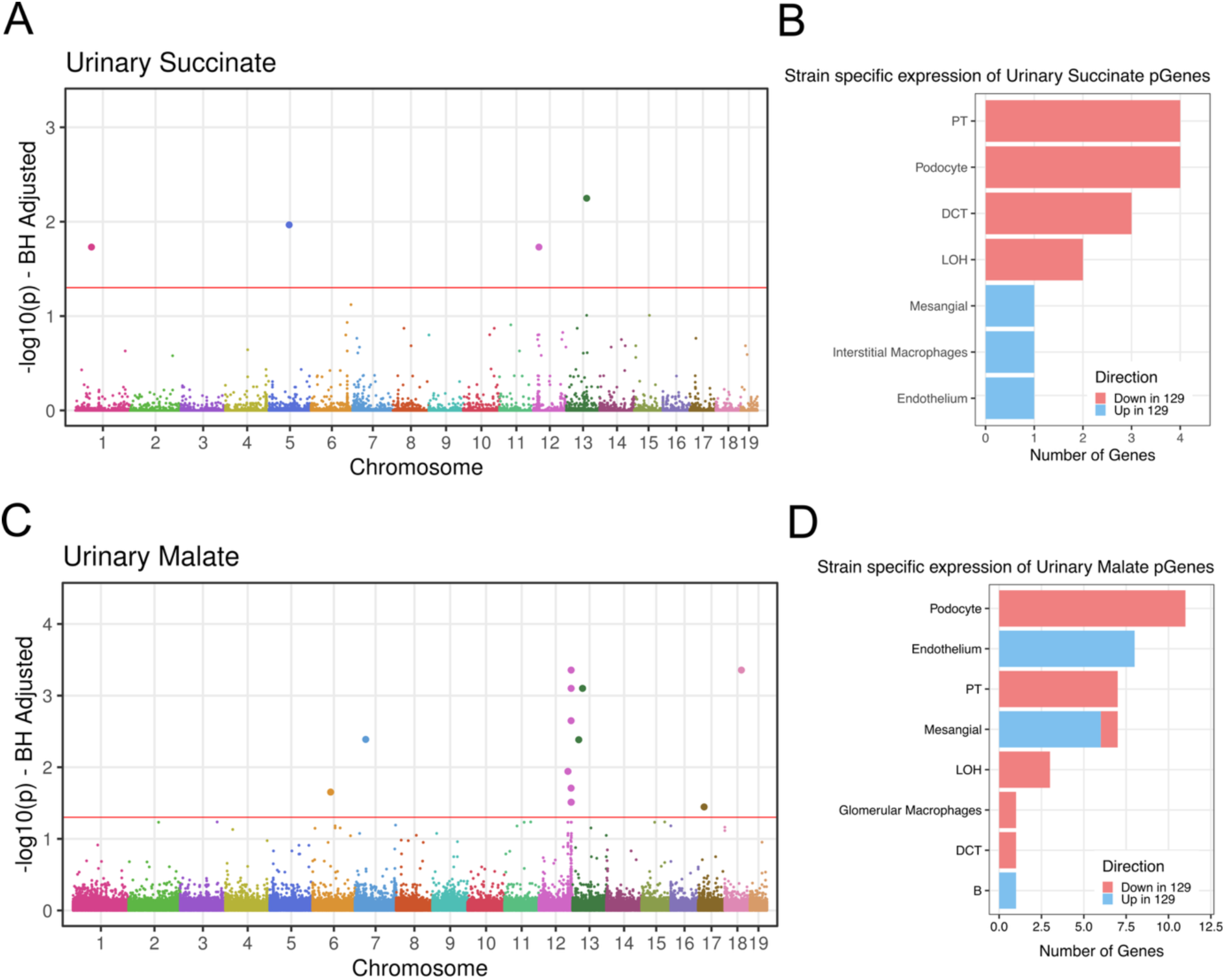
QTL mapping and cell-type-specific expression of urinary organic acid pGenes in the F2 cohort. (A) Manhattan plot showing the results of linear mixed model (LMM) QTL mapping for urinary succinate levels in the F2 cohort. (B) Bar plot showing the number of urinary succinate pGenes with strain-specific differential expression across kidney cell types, stratified by direction of effect (red, lower in 129AR; blue, higher in 129AR, relative to BL6AR). (C) Manhattan plot showing LMM QTL mapping results for urinary malate levels. (D) Bar plot showing strain-specific differential expression of urinary malate pGenes across kidney cell types, stratified by direction of effect. In panels A and C, each point represents a genetic locus, with chromosomal position on the x-axis and BH-adjusted -log10(p-value) on the y-axis. The horizontal red line indicates the significance threshold (BH-adjusted p < 0.05). Significant loci are enlarged. Cell type abbreviations: PT, proximal tubule; DCT, distal convoluted tubule; LOH, loop of Henle.

**Extended Figure 4.**
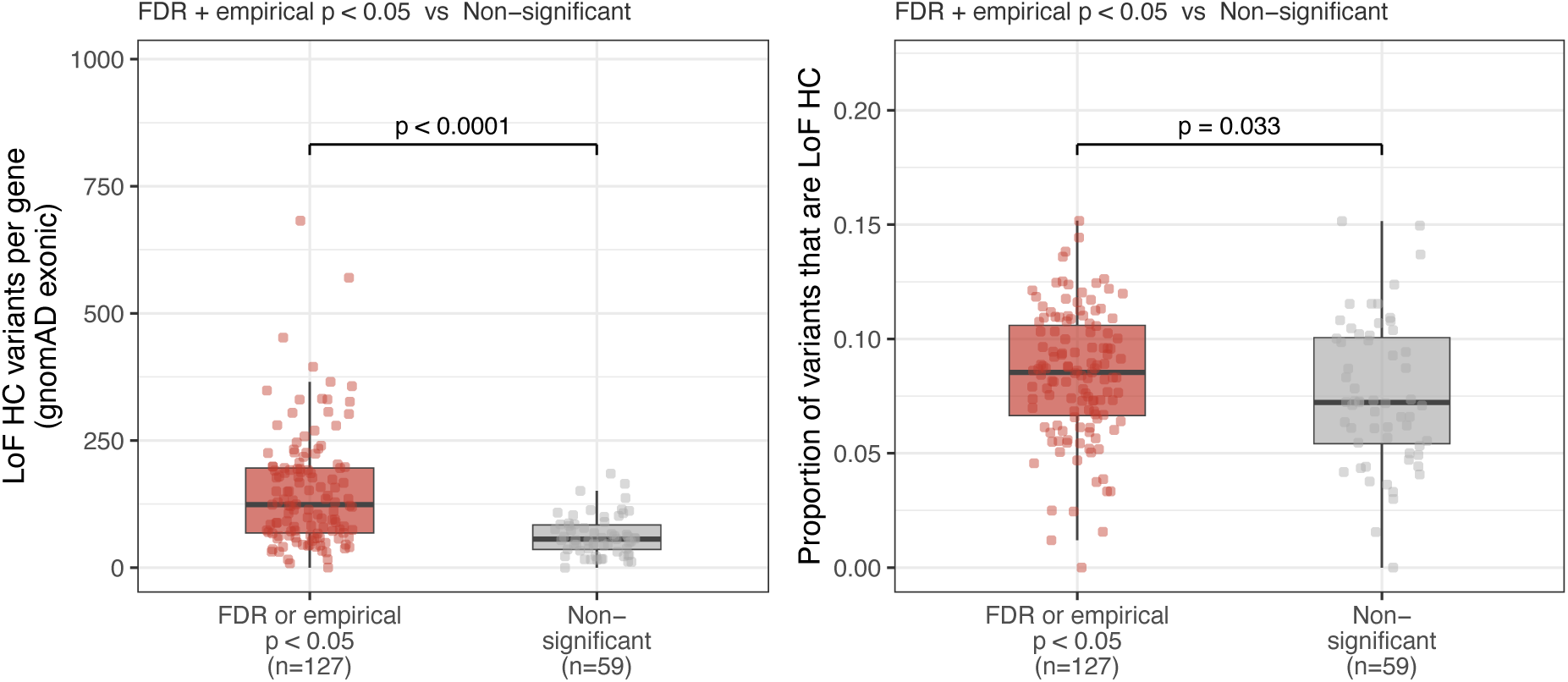
Significant albuminuria pGenes are enriched for high-confidence loss-of-function variants. Left: Boxplot showing the number of gnomAD exonic high-confidence (HC) loss-of-function (LoF) variants per gene in albuminuria pGenes reaching FDR or empirical significance (FDR or empirical p < 0.05, n=127) versus non-significant pGenes (n=59). Right: Boxplot showing the proportion of variants that are HC LoF variants per gene across the same groups. Statistical comparisons were performed using the Wilcoxon rank-sum test. p-values are shown above each comparison.

**Extended Figure 5.**
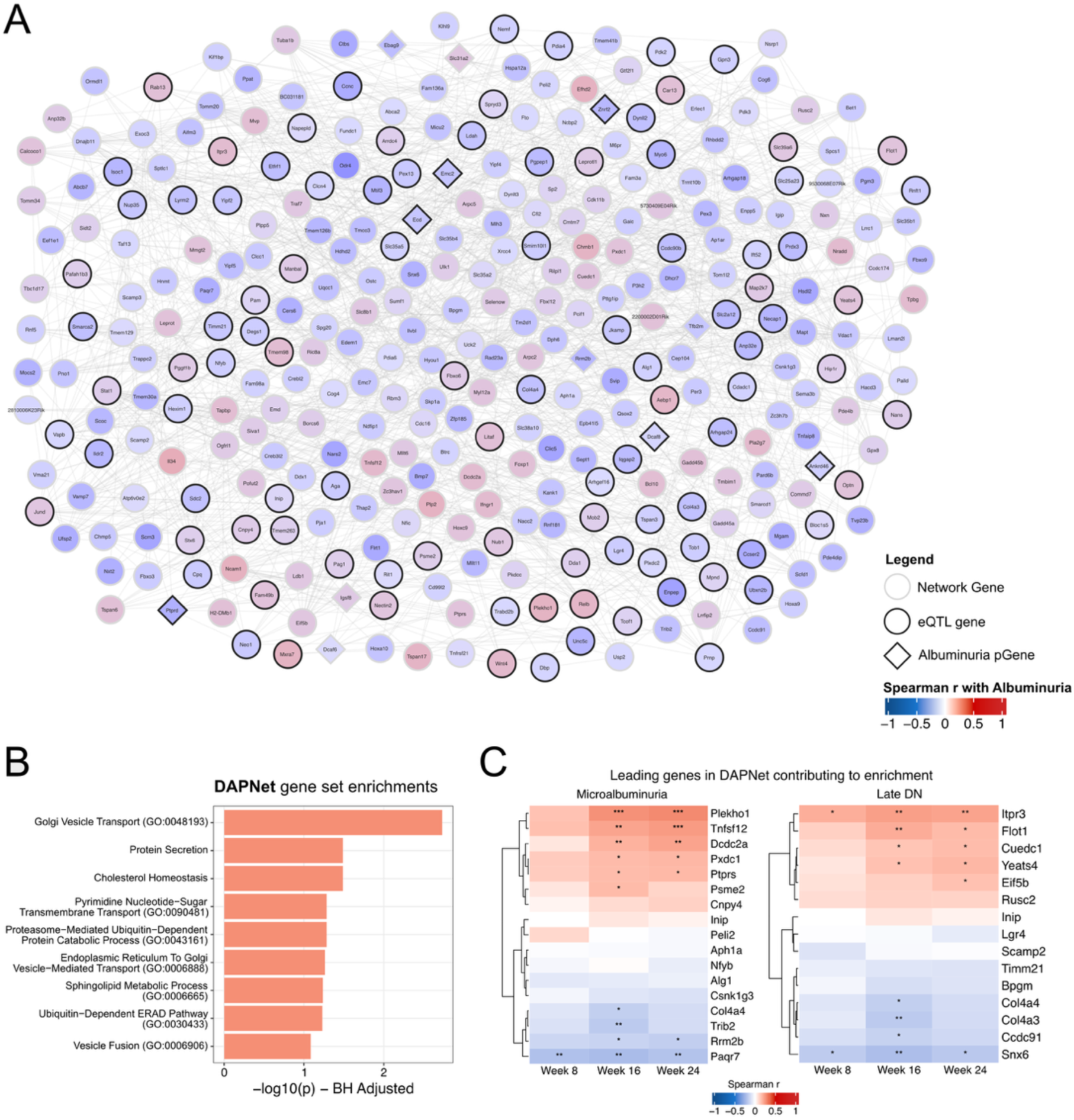
Disease Associated Podocyte Network (DAPNet) identified from the unbiased analysis of gene co-expression networks in podocytes. (A) Network visualization of DAPNet genes. Nodes represent genes; edges represent protein-protein interactions from STRING with a confidence score greater than 0.5. Node shape indicates gene category: circles denote network genes, and diamonds denote genes that are also albuminuria pGenes. Nodes outlined in black are additionally identified as eQTL genes. Node color reflects the Spearman correlation of each gene with albuminuria at 24 weeks, ranging from blue (strong negative correlation, r = -1) to red (strong positive correlation, r = 1). The full list of DAPNet genes is provided in Supplementary Table 16. (B) Gene set enrichment analysis of DAPNet genes. Bars show BH-adjusted -log10(p) values for significantly enriched Gene Ontology biological process terms. Top enrichments include Golgi vesicle transport, protein secretion, cholesterol homeostasis, and sphingolipid metabolic process, reflecting the predominant roles of membrane trafficking and lipid metabolism within the network. (C) Leading-edge genes driving DAPNet enrichment for human diabetic nephropathy GWAS signals at the microalbuminuria (left) and late DN (right) stages. Heatmaps display Spearman correlation coefficients between each gene and albuminuria at weeks 8, 16, and 24. Asterisks denote statistical significance (* p < 0.05, ** p < 0.01, *** p < 0.001).

**Extended Figure 6.**
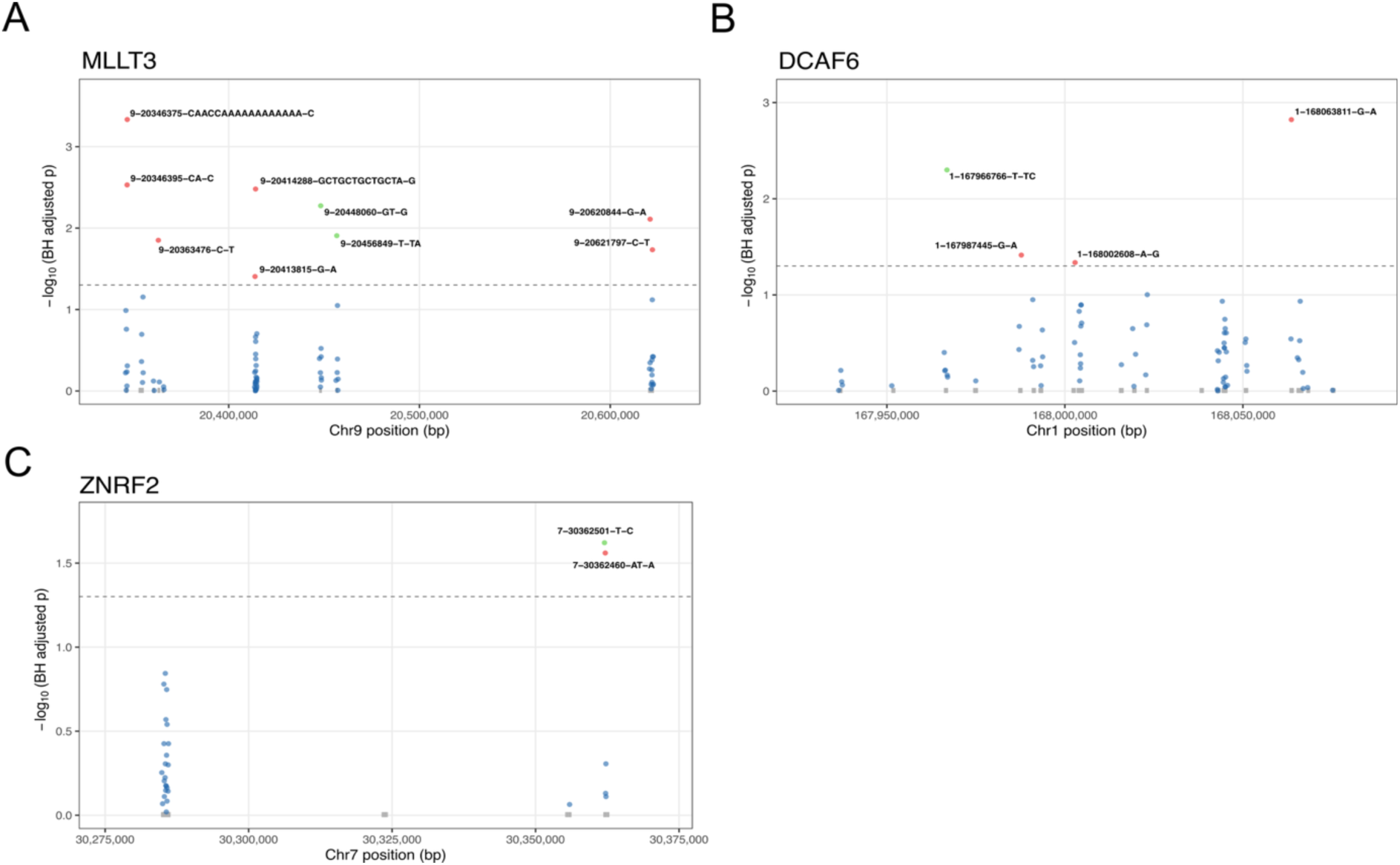
UACR-associated variants in functionally validated DAPNet candidate genes. (A) Locus plot showing variants within the MLLT3 locus on chromosome 9 associated with urine albumin-to-creatinine ratio (UACR) in diabetic participants of the All of Us Research Program. Red points indicate adverse variants and green points indicate protective variants exceeding the significance threshold (dashed line, BH-adjusted p < 0.05). (B) Locus plot for the DCAF6 locus on chromosome 1. (C) Locus plot for the ZNRF2 locus on chromosome 7. In all panels, each point represents a single variant, with chromosomal position on the x-axis and BH-adjusted association p-value on the y-axis. Labelled variants exceed the significance threshold. Blue points are non-significant variants. Association testing was performed using a linear mixed model accounting for age, sex, race, and fasting glucose levels, with gene-level FDR correction applied across all tested pGene orthologs.

## Supplementary Section

## Supplementary Tables

**Supplementary Table 1.** Histological scoring criteria for kidney pathology assessment in F2 mice. Semi-quantitative scoring system used to evaluate renal damage, adapted from criteria for human diabetic nephropathy. Features scored include mesangial expansion, nodular sclerosis, global glomerulosclerosis, tubular basement membrane thickening, interstitial fibrosis and tubular atrophy, and arteriolar hyalinosis. Scores for each feature and their corresponding definitions are shown.

**Supplementary Table 2.** Phenotypic data for all F2 mice (n=279). Includes histological scores, urinary organic acid metabolite levels at 24 weeks (µmoles/24 hours), blood glucose levels at 8, 16, and 24 weeks, 24-hour urine volume, albumin concentration, urinary albumin excretion, and normalised CD68+ area.

**Supplementary Table 3.** Significant pQTLs for urinary albumin excretion (UAE) identified by linear mixed model mapping in the F2 cohort. Each row represents a lead SNP for a significant locus after Benjamini-Hochberg correction (adjusted p < 0.05). Columns include SNP identifier, chromosome, haplotype block size, block coordinates, number of genes in block, minor allele frequency, alleles, raw and adjusted p-values, and effect size (beta) at 8, 16, and 24 weeks.

**Supplementary Table 4.** Albuminuria pGenes: 192 candidate genes prioritised from UAE pQTL loci. Genes were retained if they harboured sequence differences between 129AR and BL6AR parental strains, had human orthologs, and were expressed in the mouse kidney. Columns include mouse gene name, originating lead SNP, cis-eQTL support (Yes/No), human ortholog, Drosophila ortholog, haplotype block size, kidney cell types expressing the gene, and known functions.

**Supplementary Table 5.** Significant pQTLs for CD68+ macrophage infiltration identified by linear mixed model mapping in the F2 cohort (adjusted p < 0.05). Columns include SNP identifier, chromosome, haplotype block coordinates and size, minor allele frequency, alleles, raw and adjusted p-values, and effect size.

**Supplementary Table 6.** CD68 pGenes: 36 candidate genes prioritised from macrophage infiltration pQTL loci after filtering for parental strain sequence differences, human orthologs, and kidney expression. Columns as in Supplementary Table 4.

**Supplementary Table 7.** Significant pQTLs for global glomerulosclerosis identified using a zero-inflated gamma regression model with permutation-based empirical p-values. Columns include SNP identifier, chromosome, haplotype block coordinates and size, minor allele frequency, alleles, raw and empirical p-values, and effect size.

**Supplementary Table 8.** Glomerulosclerosis pGenes: 25 candidate genes prioritised from glomerulosclerosis pQTL loci after filtering for parental strain sequence differences, human orthologs, and kidney expression. Columns as in Supplementary Table 4.

**Supplementary Table 9.** Significant pQTLs for urinary succinate levels at 24 weeks identified by linear mixed model mapping (adjusted p < 0.05). Columns include SNP identifier, chromosome, haplotype block coordinates and size, allele frequency, minor allele frequency, alleles, raw and adjusted p-values, and effect size.

**Supplementary Table 10.** Succinate pGenes: 12 candidate genes prioritised from urinary succinate pQTL loci after filtering for parental strain sequence differences, human orthologs, and kidney expression. Columns as in Supplementary Table 4.

**Supplementary Table 11.** Significant pQTLs for urinary malate levels at 24 weeks identified by linear mixed model mapping (adjusted p < 0.05). Columns include SNP identifier, chromosome, haplotype block coordinates and size, minor allele frequency, alleles, raw and adjusted p-values, and effect size.

**Supplementary Table 12.** Malate pGenes: 27 candidate genes prioritised from urinary malate pQTL loci after filtering for parental strain sequence differences, human orthologs, and kidney expression. Columns as in Supplementary Table 4.

**Supplementary Table 13.** Significant cis-eQTLs identified in whole kidney RNA-seq data from the F2 cohort (n=279). A total of 5,494 cis-eGenes were identified at FDR < 0.05 using the Benjamini-Hochberg procedure. Cis-eQTLs were defined as SNP-gene pairs on the same chromosome within ±1 Mb of the gene midpoint. Columns include gene name, lead SNP, gene chromosome and position, distance from SNP, alleles, minor allele frequency, effect size (beta), standard error, nominal p-value, and FDR q-value.

**Supplementary Table 14.** Significant trans-eQTLs identified in whole kidney RNA-seq data from the F2 cohort (n=279). A total of 50 trans-eGenes were identified after Bonferroni correction across the full trans search space. Trans-eQTLs were defined as SNP-gene pairs on different chromosomes or beyond ±1 Mb of the gene midpoint. Columns include gene name, lead SNP, gene chromosome and position, distance from SNP, alleles, minor allele frequency, effect size (beta), standard error, and nominal p-value.

**Supplementary Table 15.** Albuminuria pGenes with prior associations to CKD or DN-related traits. Of the 192 albuminuria pGenes, 52 had previously reported associations with CKD, type 1 or type 2 diabetes, eGFR decline, or hypertension in human GWAS. Columns include mouse gene name, associated human traits, GWAS disease trait descriptions, minimum reported p-value, associated SNPs, study accession numbers, and PubMed IDs.

**Supplementary Table 16.** Summary of 95 cell-type-specific gene coexpression networks constructed by WGCNA from parental strain single-cell RNA-seq data. For each network, columns include network name, cell type of origin, network size, Spearman correlation with albuminuria at each timepoint with adjusted p-values, overlap with albuminuria pGenes, number of differentially expressed genes between 129 and BL6 strains, number of cis- and trans-eGenes within the network, and MAGMA enrichment p-values for microalbuminuria, macroalbuminuria, and ESRD from the van Zuydam et al. 2018 DN GWAS.

**Supplementary Table 17.** Drosophila RNAi lines used for nephrocyte knockdown experiments. Lists the fly line name and Bloomington Drosophila Stock Center reference number for each UAS-RNAi line used in the filtration and actin imaging assays.

---

**Supplementary Figure 1.**
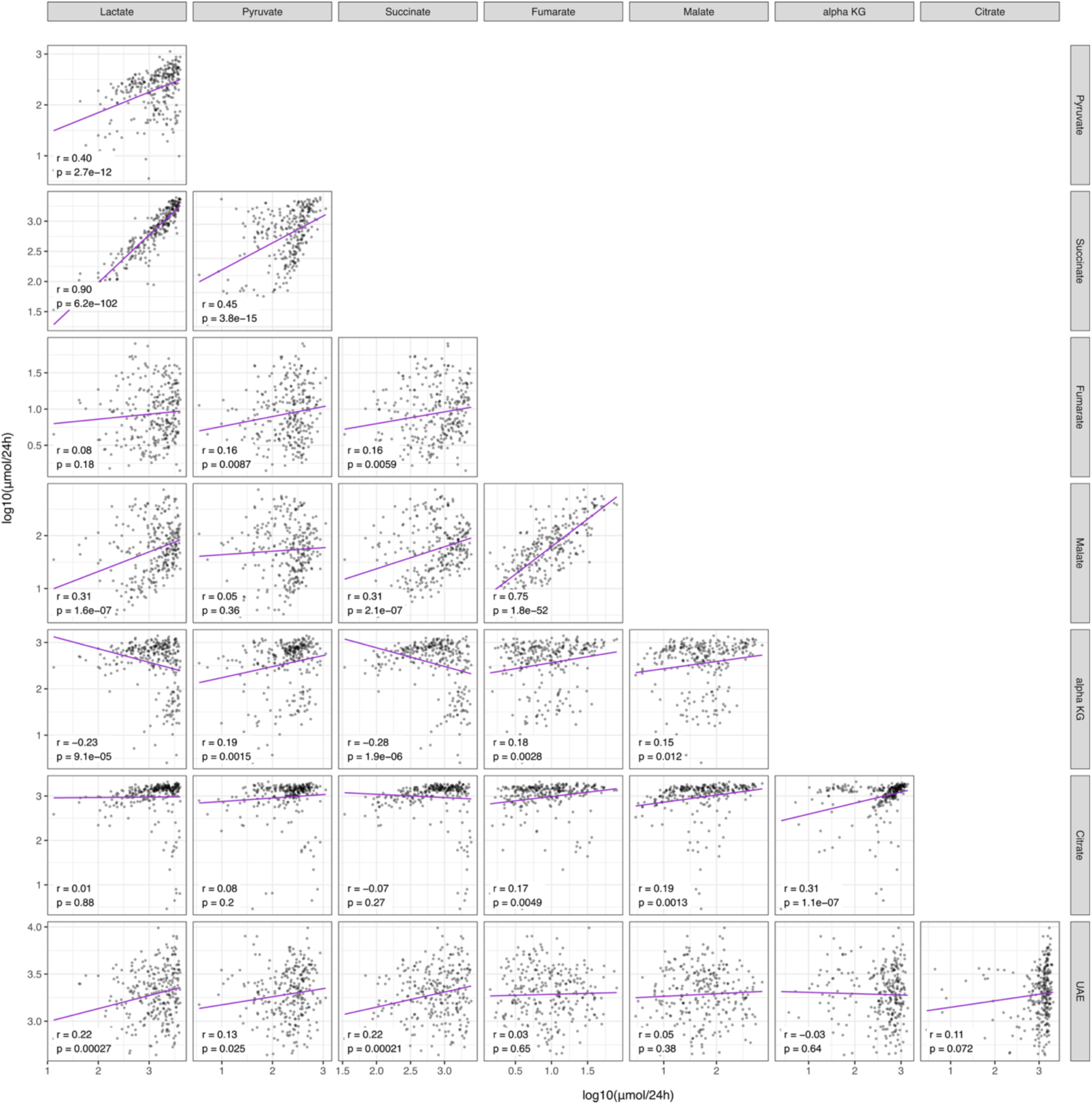
Urinary metabolites correlate with albuminuria in the F2 cohort. Pairwise Spearman correlation plots showing relationships among urinary organic acid metabolites (lactate, pyruvate, succinate, fumarate, malate, alpha-KG, and citrate) and urinary albumin excretion (UAE) at 24 weeks of age. Each panel shows log-transformed metabolite concentrations on the x- and y-axes, with a linear regression line overlaid in purple. Spearman correlation coefficient (r) and associated p-value are shown in each panel. UAE is shown on the y-axis of the bottom row. Significant correlations (p < 0.05) indicate co-variation among TCA cycle intermediates and their association with albuminuria severity.

**Supplementary Figure 2.**
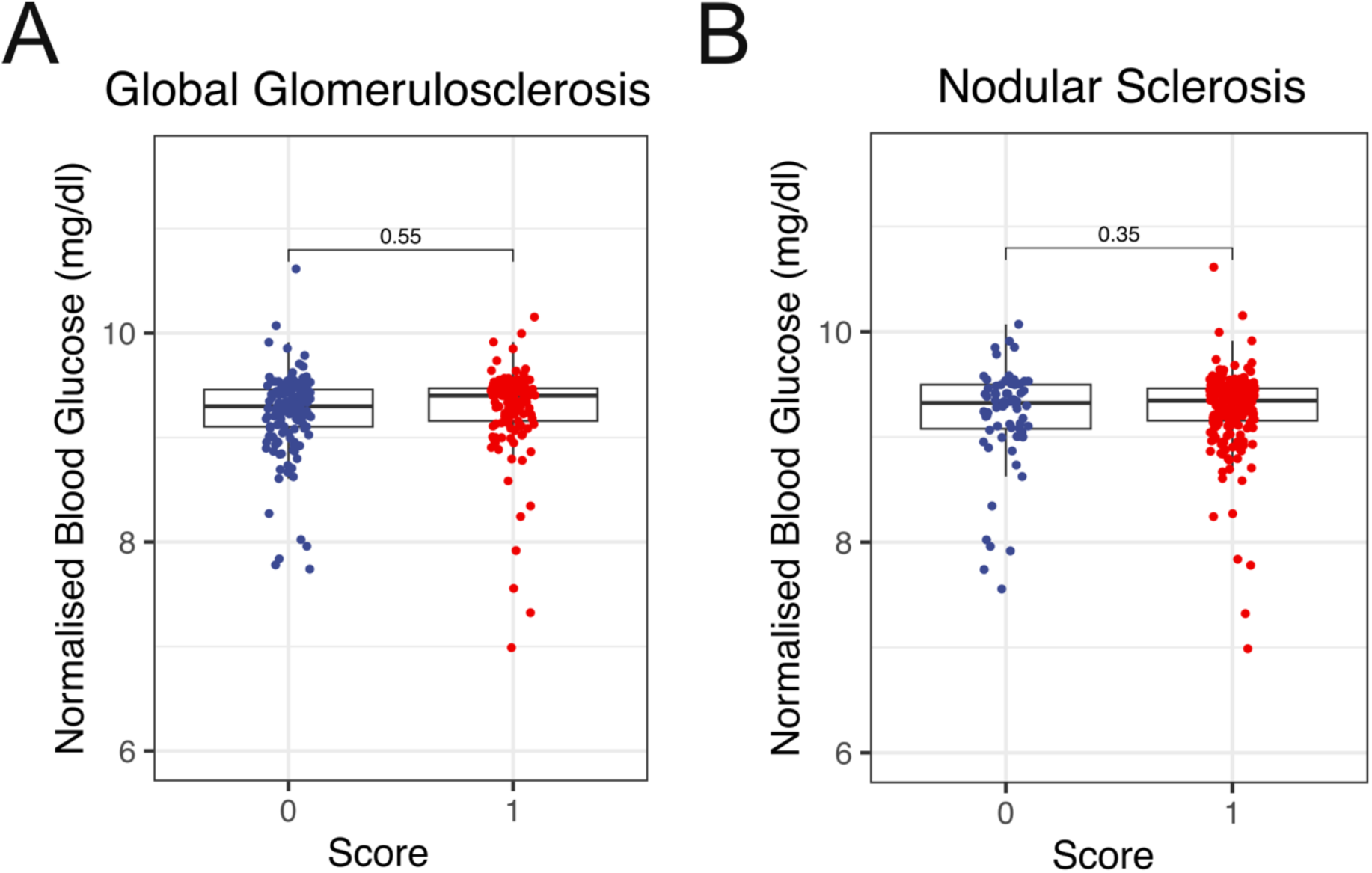
Blood glucose levels are comparable across histological severity groups in the F2 cohort. Boxplots showing log-normalised blood glucose levels stratified by (A) global glomerulosclerosis score and (B) nodular sclerosis score in F2 mice at 24 weeks. Score 0 indicates absence and score 1 indicates presence of the respective lesion. Each point represents an individual F2 mouse. Statistical comparisons were performed using a two-tailed Student’s t-test, with p-values shown above each comparison.

**Supplementary Figure 3.**
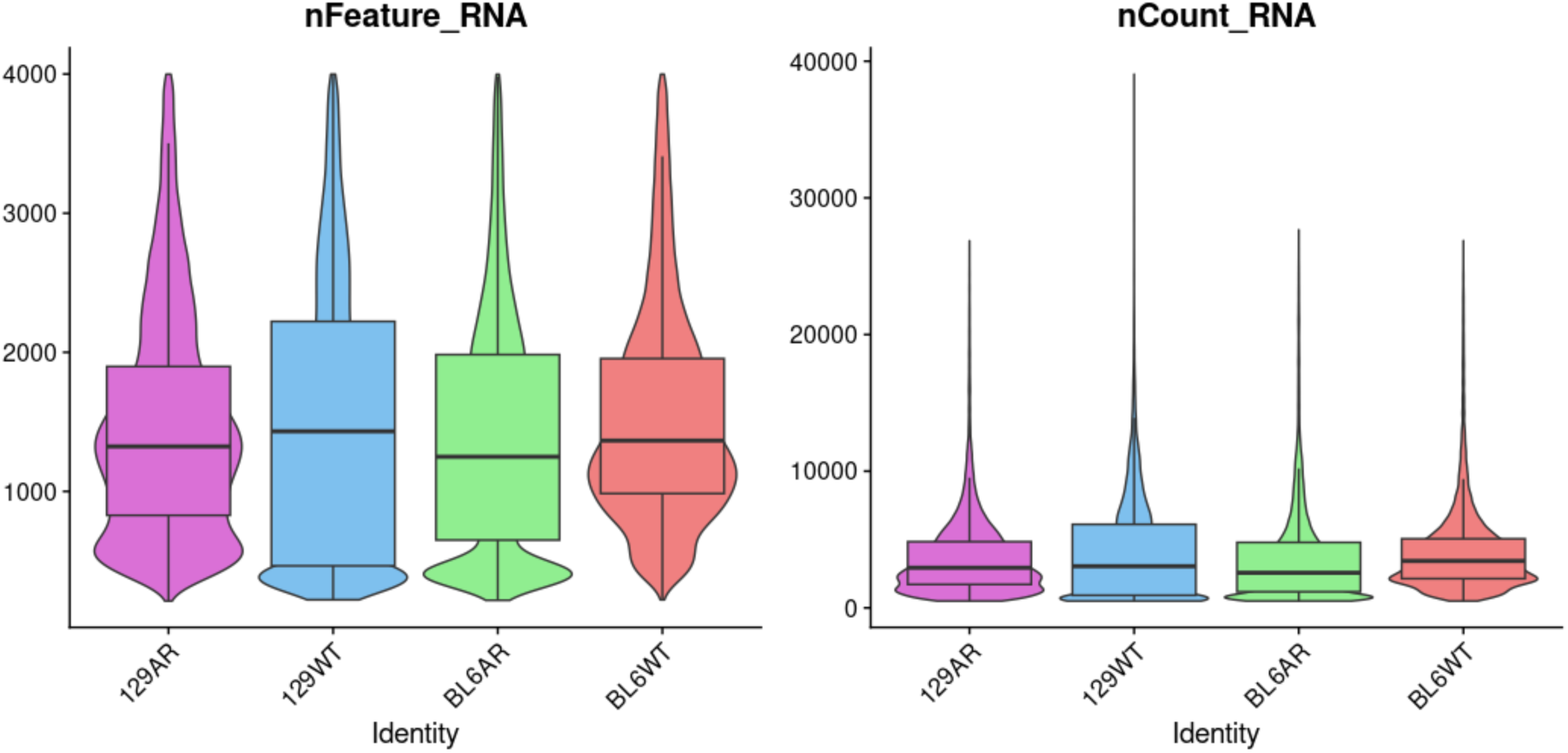
Violin plots showing the QC metrics for the single cell dataset. The panel on the left shows the number of unique genes captured for each mouse strain, denoted by “nFeature_RNA”. The panel on the right shows the number of unique molecular identifiers (UMIs) captured for each strain, as given by “nCount_RNA”. Both panels show a similar capture rate across all strains.

**Supplementary Figure 4.**
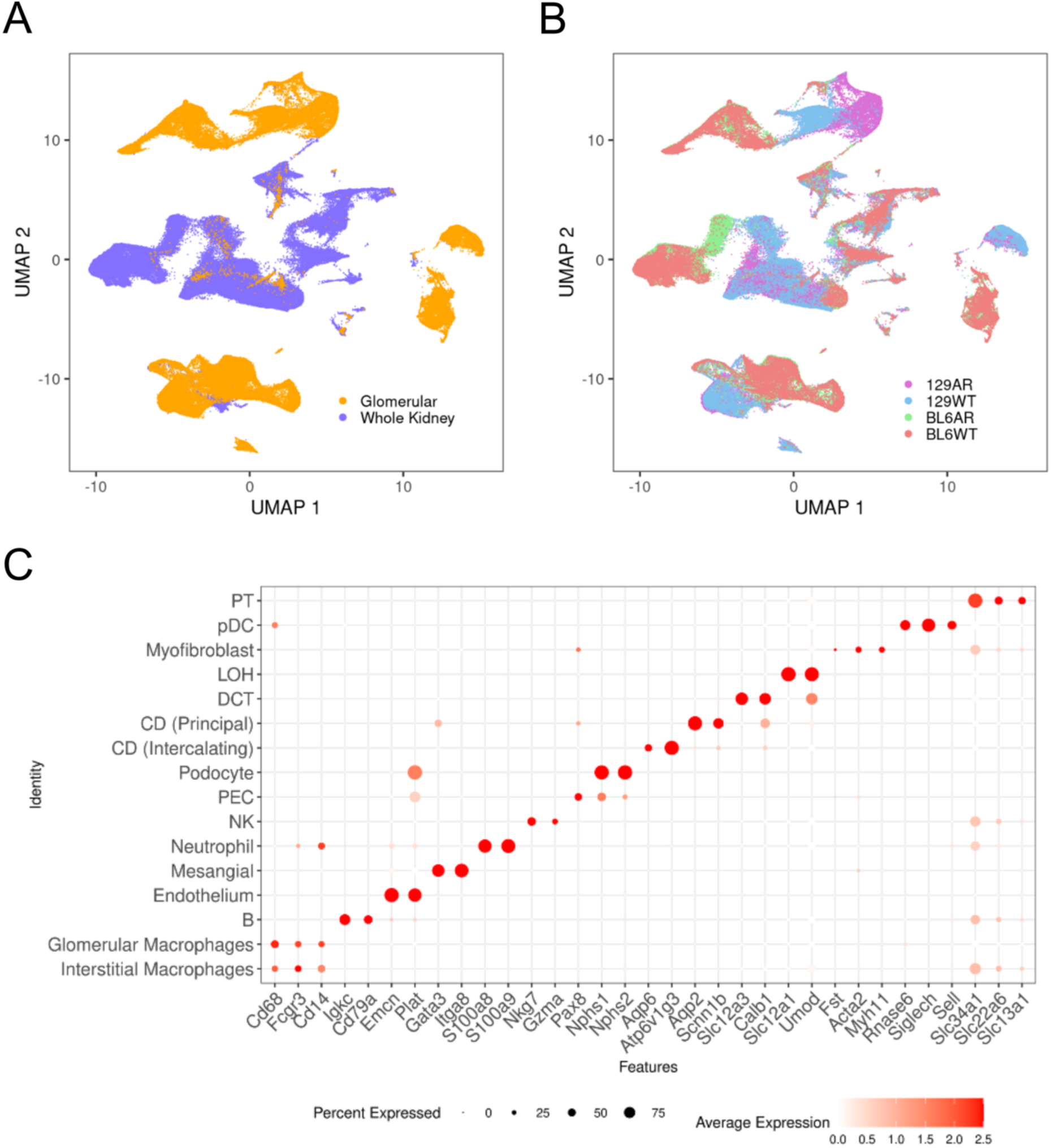
Single cell plots describing the cells captured in this dataset. (A) and (B) are UMAP plots coloring cells by cell extract (A) or by mouse strain (B). (C) shows a dotplot coloring cells for various marker genes used to annotate the clusters after unsupervised clustering and dimensionality reduction.

**Supplementary Figure 5.**
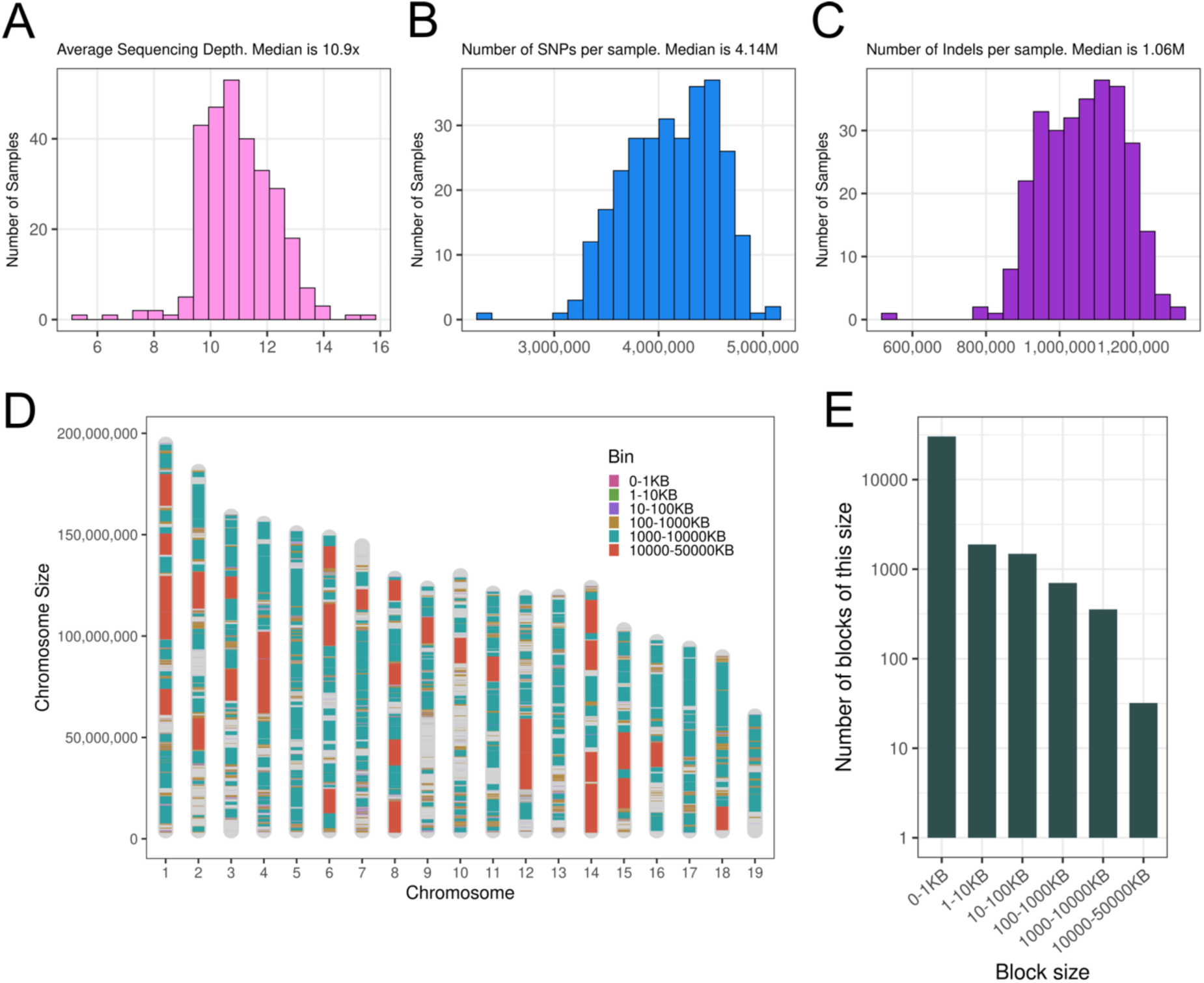
Whole-genome sequencing metrics for the F2 cohort. Histograms showing the distribution of (A) average sequencing depth (median 10.9x), (B) the number of single-nucleotide polymorphisms (SNPs) per sample (median 4.14M), and (C) the number of insertions and deletions (indels) per sample (median 1.06M) across the F2 cohort. (D) Chromosomal visualization of haplotype blocks constructed using the D’ linkage disequilibrium metric, with blocks colour-coded by size category. (E) Bar plot showing the distribution of haplotype block counts across size bins, illustrating the range from short blocks under 1 kb to large blocks exceeding 1 Mb.

